# The Habitat Amount Hypothesis implies negative effects of habitat fragmentation on species richness and occurrence

**DOI:** 10.1101/2020.02.02.930784

**Authors:** Santiago Saura

## Abstract

The Habitat Amount Hypothesis (HAH) predicts that species richness, abundance or occurrence in a habitat site increases with the amount of habitat in the ‘local landscape’ defined by an appropriate distance around the site, with no distinct effects of the size of the habitat patch in which the site is located. It has been stated that a consequence of the HAH, if supported, would be that it is unnecessary to consider habitat configuration to predict or manage biodiversity patterns, and that conservation strategies should focus on habitat amount regardless of fragmentation. Here, I assume that the HAH holds and apply the HAH predictions to all habitat sites over entire landscapes that have the same amount of habitat but differ in habitat configuration. By doing so, I show that the HAH actually implies clearly negative effects of habitat fragmentation, and of other spatial configuration changes, on species richness, abundance or occurrence in all or many of the habitat sites in the landscape, and that these habitat configuration effects are distinct from those of habitat amount in the landscape. I further show that, contrary to current interpretations, the HAH is compatible with a steeper slope of the species-area relationship for fragmented than for continuous habitat, and with higher species richness or abundance for a single large patch than for several small patches with the same total area (SLOSS). This suggests the need to revise the ways in which the HAH has been interpreted and can be actually tested. The misinterpretation of the HAH has arisen from confounding and overlooking the differences in the spatial scales involved: the individual habitat site at which the HAH gives predictions, the local landscape around an individual site, and the landscapes or regions (with multiple habitat sites and different local landscapes) that need to be analysed and managed. The HAH has been erroneously viewed as negating or diminishing the relevance of fragmentation effects, while it actually supports the importance of habitat configuration for biodiversity. I conclude that, even in the cases where the HAH holds, habitat fragmentation and configuration are important for understanding and managing species distributions in the landscape.

## 1. THE HABITAT AMOUNT HYPOTHESIS (HAH)

Understanding how habitat amount and configuration affect species richness, occurrence or abundance has been one of the major focuses of research in ecology and biogeography, given its central importance for conservation planning and for landscape (or seascape) management. In particular, there is a long-standing and vigorous debate on the relative importance of habitat amount (total area of habitat) and habitat spatial configuration (the spatial arrangement of the habitat) for biodiversity patterns and persistence (Wilcox & Murphy, 1985; Fahrig, 2013; Fahrig, 2015; Haddad et al., 2015; Hanski, 2015; Martin, 2018). The most widely accepted conceptual model, and the predominant consensus among ecologists, has been that both habitat amount and configuration (e.g. fragmentation) matter and need to be considered in conservation management (e.g. Haddad et al., 2015; Hanski, 2015).

Fahrig (2013) challenged this conceptual model by proposing the Habitat Amount Hypothesis (HAH). The HAH states that species richness, occurrence or abundance in a given habitat site should increase with the amount of habitat in the circular ‘local landscape’ defined by a certain distance *D* surrounding that site (scale of effect), with no additional (distinct) effects of the area of the habitat patch in which the habitat site is located. Although in the presentation and discussion of the HAH Fahrig (2013) focused on species richness, she explicitly noted that the HAH could equally be applied to the abundance or probability of occurrence of individual species. Indeed, because species richness is the sum of the occurrence of individual species, that the HAH holds for species richness means that it should also hold for the occurrence of at least most of the individual species (Fahrig, 2013). Ultimately, it may be more valid to test the hypothesis by accumulating tests of the HAH across many species, than by conducting tests on species richness (Fahrig 2013; Fahrig, 2015). Therefore, I here consider that the response variable predicted through the HAH can be either species richness or the abundance or occurrence of an individual species (depending on the focus and interest of a particular study), using in all cases the same explanatory variable (amount of habitat in the local landscape around each site).

The implications of the HAH are important, and the interpretations that have been made of this hypothesis since it was proposed are strong, both from a conceptual and conservation point of view. Fahrig (2013) stated that if the HAH was supported, considering habitat configuration independent of habitat amount would be unnecessary; there would be no distinct effects of habitat patch size and isolation on species richness, abundance or occurrence. Torrenta and Villard (2017) stated that the HAH challenges the contention that fragmentation effects in the strict sense actually exist, which has important implications, both from a theoretical and a conservation perspective. Bueno and Peres (2019) stated that support for the HAH would imply that biodiversity conservation strategies should focus on retaining the maximum overall amount of habitat regardless of its configuration. MacDonald, Anderson, Acorn and Nielsen (2018) stated that the HAH “negates fragmentation effects and predicts that only total habitat area determines the diversity of species persisting on fragmented landscapes”. Haddad et al. (2017) stated that “if supported, the Habitat Amount Hypothesis would fundamentally transform how landscapes are conserved by shifting focus away from considerations of habitat configuration and connectivity, toward total habitat area alone”. Melo, Sponchiado, Cáceres and Fahrig (2017) concluded, after finding support for the HAH, that their results “support the notion that biodiversity protection policies should focus on habitat amount, irrespective of its spatial configuration”. Since it was proposed, the HAH has been supported by some studies (e.g. Melo et al., 2017; Rabelo, Bicca-Marques, Aragón, & Nelson, 2017; Seibold et al., 2017) but rejected by others (e.g. Evju & Sverdrup-Thygeson, 2016; Haddad et al., 2017; Lindgren & Cousins, 2017); see Martin (2018) for a recent meta-analysis.

I here challenge the interpretation that the HAH implies that habitat fragmentation or configuration is not important for species richness, abundance or occurrence. I argue that even if the HAH holds, conservation strategies cannot focus only on the amount of habitat in the landscape, and should also consider the spatial arrangement of habitat in the landscape. I do not provide any arguments or empirical results supporting or rejecting the HAH. I assume that the HAH actually holds. I elaborate on the implications that the HAH actually has, assuming that this hypothesis holds, for the distribution of species richness or occurrence in a landscape or region, which I illustrate with a variety of habitat spatial configurations to which I apply the predictions of the HAH. By doing so, I show that the HAH actually implies clearly negative effects of habitat fragmentation on species richness, abundance or occurrence in a landscape or region, contrary to prevailing understanding on the implications of the HAH. I explain that the misinterpretation of the implications of the HAH in previous studies has been due to the confusion of different spatial scales: the individual habitat site, the local landscape around that site, and the scale of an entire landscape or region comprising multiple habitat sites and local landscapes around them. Not clearly differentiating these spatial scales has led to misunderstandings on what the HAH really implies and how it can be actually tested.

## 2. THE HABITAT AMOUNT HYPOTHESIS IMPLIES THE IMPORTANCE OF HABITAT FRAGMENTATION AND CONFIGURATION FOR BIODIVERSITY

### 2.1. The local landscape and the habitat sites in the HAH

A key specification of the Habitat Amount Hypothesis (HAH) is the distance *D* that defines the circular ‘local landscape’ (scale of effect) over which the amount of habitat determines the response variable (usually species richness, abundance or occurrence) in a site (Fahrig, 2013). The distance *D* has no universal value, but has to be determined empirically in each case as the distance at which the habitat amount shows the strongest correlation with the response variable. Although *D* will be different in each case, the studies that have so far evaluated the HAH have found *D* to range from 40 m to 5500 m (Evju & Sverdrup-Thygeson, 2016; Lindgren & Cousins, 2017; Melo et al., 2017; Rabelo et al., 2017; Seibold et al., 2017; Torrenta & Villard, 2017; Gardiner et al., 2018; Thiele, Kellner, Buchholz, & Schirmel, 2018; Vieira, Almeida-Gomes, Delciellos, Cerqueira, & Crouzeilles, 2018; Bueno & Peres, 2019), with an average *D* value of about 1000 m. I make no assumption on the particular value of *D*, but just assume that the HAH holds, and therefore, that such scale of effect exists and has been determined in each particular case.

The habitat sites in which species are sampled in the HAH are equally sized and are much smaller than the local landscape. Obviously, the HAH would be of no use if habitat sites were of the same or similar size as the local landscape; all sites would have, in such an extreme case, the same amount of habitat in the local landscape and hence the same value of the response variable as predicted by the HAH. Although no specific indication is given on the relative sizes of the sites and the local landscapes in the HAH, the literature on the HAH has assessed local landscapes with an area that varies from a few dozen times to several tens of thousand times larger than the area of the habitat sites in which the species were sampled (Evju & Sverdrup-Thygeson, 2016; Lindgren & Cousins, 2017; Melo et al., 2017; Seibold et al., 2017; Torrenta & Villard, 2017; Thiele et al., 2018; Vieira et al., 2018). Hereafter, I just assume that the habitat sites are equally sized and much smaller than the local landscape; the particular relative size of the two (site vs local landscape) is not important for the illustrative cases shown and the arguments made hereafter (but see Appendix S1 in Supporting Information for details).

I will here use, for brevity, the term ‘landscape’ to refer to an area of any extent in which habitat is distributed. It may or may not be larger than the extent of a ‘local landscape’ (circle with radius *D*) as defined in the HAH; this will be specified in each particular case only when necessary. There is a key distinction between these two terms (‘landscape’ and ‘local landscape’). When I refer to a ‘landscape’, I am concerned with the response variable (e.g. species richness) in *all the habitat sites* found within the landscape; see for example any of the eight rectangular landscapes in Figure 1. When I refer to a ‘local landscape’, I take the HAH perspective and focus in an *individual habitat site* located in the centre of *that local landscape*; see any of the examples of individual habitat sites and their local landscapes highlighted in green within the landscapes of Figure 1. In this latter case, the concern is the response variable (e.g. species richness) only in a focal individual habitat site; this is the one for which the HAH gives the predictions based on the habitat amount in the local landscape around that site. Other habitat sites different from the focal habitat site may sum to the total amount of habitat in the local landscape (if the sites are close enough to the focal one) but the HAH does not predict the response variable in the *rest of the habitat sites* from the information of *that local landscape*. In the HAH, the local landscape is determined specifically for a single focal site, and the HAH uses the information in that local landscape to give predictions for that focal habitat site only.

**FIGURE 1.**
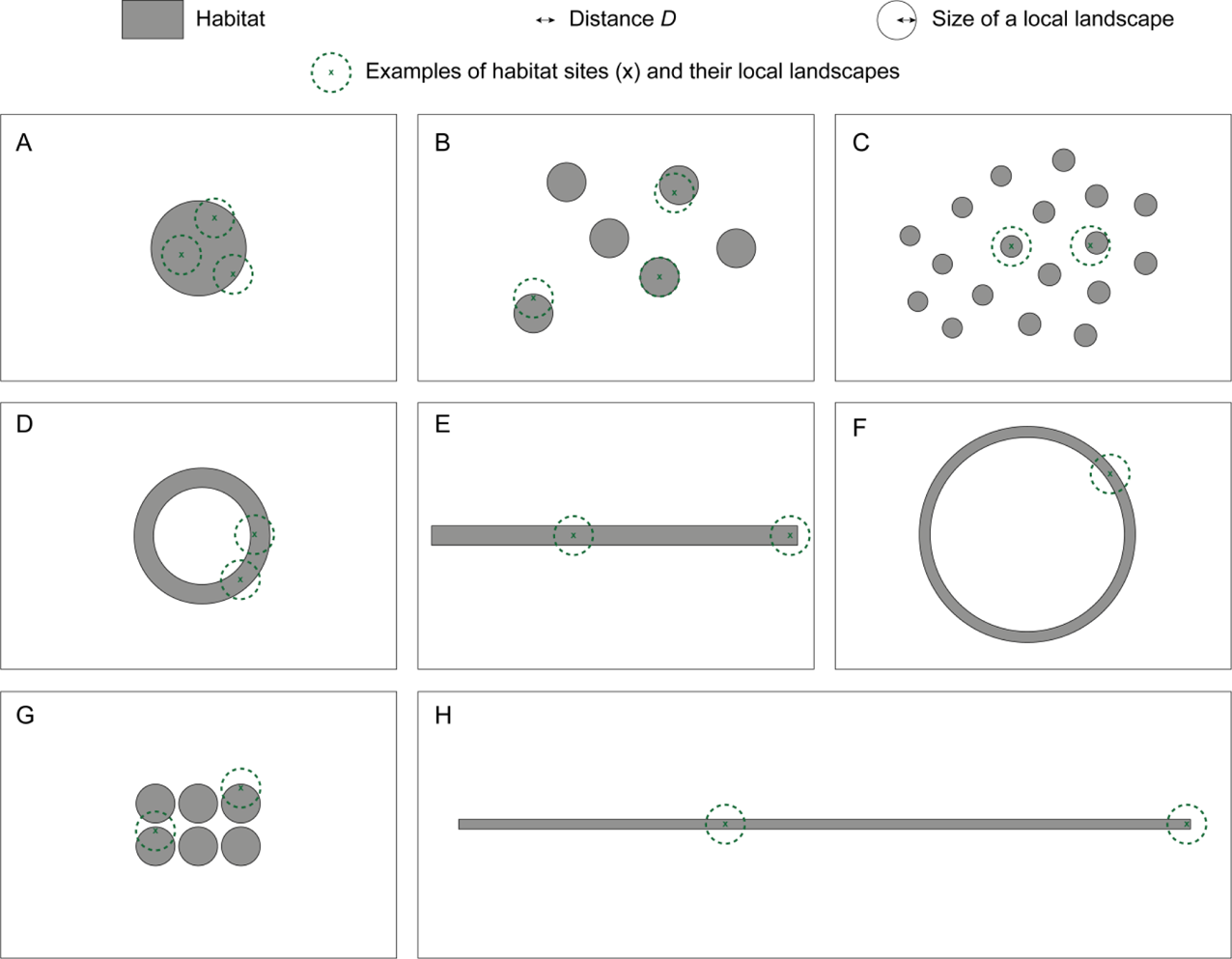
Eight landscapes with the same habitat amount but different habitat spatial configuration. The three first landscapes (A-C) correspond to an increase in habitat fragmentation, with patch size larger than (A), equal to (B) or smaller than (C) the size of a local landscape in the habitat amount hypothesis, which is given by the distance *D* (scale of effect). Landscapes D, E, F and H have a single habitat patch (as in A) but with different degrees of perforation or elongation. Landscape G is as B but with the habitat patches much closer to each other. The crosses (x) and circles in green colour indicate some examples of habitat sites and their local landscapes (circles of radius *D* around the sites) that differ in their location within landscapes A-H and in the amount of habitat in the local landscape.

I will here consider that the habitat patches, and the landscapes that contain the habitat patches, may or may not be smaller than the scale of effect (extent of the local landscape, given by *D*), which is consistent with the studies on the HAH (Appendix S2). In any case, I will show that the conclusions below hold equally for patches larger or smaller than *D*.

### 2.2. Evaluating the HAH predictions in practice: landscapes with the same habitat amount but different habitat spatial configuration

I here consider two illustrative sets of landscapes that have the same amount of habitat but differ in the spatial configuration of it (Figures 1 and 3). The landscapes in Figure 1 are larger than a local landscape in the HAH (they are larger than a circle with radius *D*), and the individual habitat patches are larger or smaller than the extent of a local landscape depending on the cases (Figure 1). The landscapes in Figure 3 have the same extent as a local landscape (they are circles with radius *D*), and hence all the individual patches contained within these landscapes are smaller than the extent of a local landscape. The habitat in each of these landscapes is distributed in equally sized habitat sites that are much smaller than *D*; being so numerous and small, they are not shown, for the sake of clarity, in Figures 1 and 3. The particular size of the habitat sites is in any case inconsequential for the purposes here (see Appendix S1 for details).

I assume that the HAH for the distance *D* holds and, thereby, I calculate, as described next in this section, the response variable (species richness, abundance or occurrence probability) in each habitat location (site) in these landscapes using solely the HAH predictions, i.e. as a function of the amount of habitat in the local landscape for each site. Some illustrative examples of habitat sites and their local landscapes are highlighted in green in Figure 1. These examples range from the full local landscape for a site covered by habitat, as in the green colour example to the left in Figure 1A, to considerably smaller percentages of habitat in the local landscape (<50%), as in the green colour examples in landscapes C and H in Figure 1.

By doing so, I obtain distributions of the response variable (e.g. species richness) that are necessarily in full accordance with the HAH, but in my elaborations and discussion I will examine the broader results for the entire landscape (considering all habitat sites together) rather than narrowly focusing on an individual habitat site only.

The formulation of the HAH does not explicitly specify or demonstrate which should be the functional form (e.g. linear or non-linear) that best relates the response variable (e.g. species richness) in the habitat site with the habitat amount in the local landscape. In the calculations presented below, I assume a linear relationship between habitat amount and the response variable, as the simplest and most parsimonious function for the illustrative cases here considered. A linear relationship is also suggested from the results of the studies on the species-area relationship that have focused on ranges of habitat areas close to those typically considered in the HAH (Appendix S3). I also assume, even if obvious, that the species response variable (species richness, abundance or occurrence) is zero in the site when there is no habitat in the local landscape. The linear relationship is hence given by:

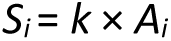

where *S_i_* is the species response variable in the habitat site *i* (species richness, abundance or occurrence in the site), *A_i_* is the amount (area) of habitat in the local landscape of the site *i* (circle of radius *D* around the centre of *i*), and *k* is a constant relating both variables.

I apply this linear relationship, individually, to each of the locations (habitat sites) in the landscapes in Figures 1 and 3, and hence obtain the HAH-predicted value of *S_i_* in each habitat site.

I then normalize the values of the species response variable as *S_norm,i_* = *S_i_* / *S_max_*, where *S_max_* is the maximum value of *S_i_* found in any of the landscapes in each of the figures in this study (a different *S_max_* value for each figure is considered). The maximum *S_max_* is found in the habitat site that, in each of these figures, has the largest habitat area in the local landscape. In Figure 1 there are some habitat patches that are larger than the extent of the local landscape (larger than a circle with radius *D*); hence some of the habitat sites have their local landscapes fully covered with habitat, and *S_max_* is equal to *k* times the size of the local landscape. In the landscapes in Figure 3 this is not however the case, and the habitat site with the maximum values of *A_i_* and *S_i_* has about one fifth of the local landscape covered by habitat; hence, for Figure 3, *S_max_* is about 0.2*k* times the size of the local landscape. The *S_norm,i_* values are independent from *k* and easier to interpret, and are hence those shown in Figures 2 and 4: *S_norm,i_*=100% corresponds to the maximum *S_i_* value found in the considered landscapes and 0% to the case of no habitat in the local landscape.

**FIGURE 2.**
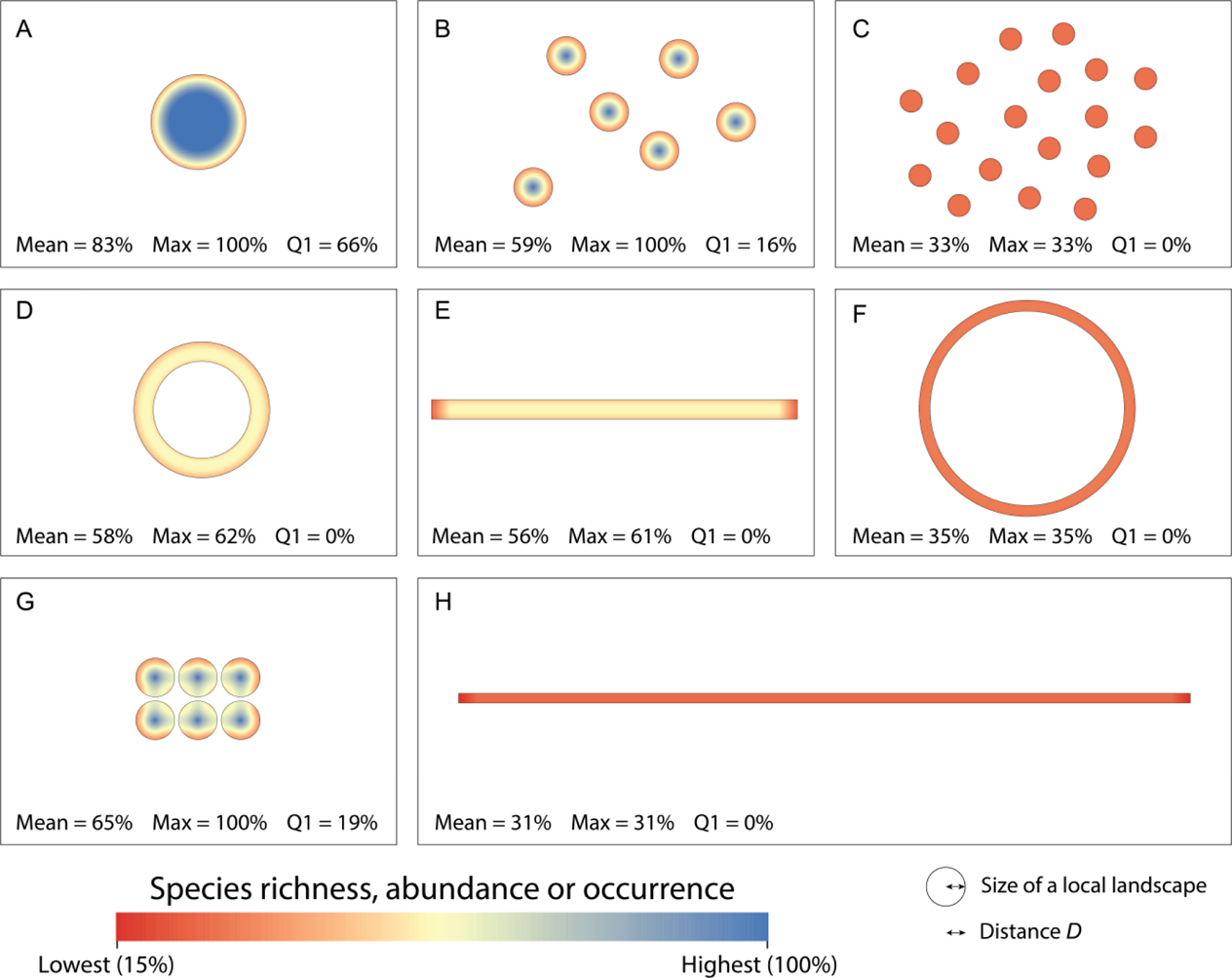
Species richness, abundance or probability of occurrence (*S_norm,i_*) given by the predictions of the Habitat Amount Hypothesis (HAH) in each of the landscapes in Figure 1, i.e. calculated for each location (habitat site) *i* as a linear function of the amount of habitat in the local landscape (distance *D*) around the site, as described in section 2.2. The figure indicates, for each landscape, the mean (Mean) and maximum (Max) value of *S_norm,i_* in the landscape as predicted by the HAH, as well as the percentage of habitat area in each landscape that has a value of *S_norm,i_* within the top quartile (Q1; within the 75-100% range). The results shows that an increased habitat fragmentation (breaking apart of habitat), elongation or perforation, while holding constant the amount of habitat, has negative effects on the species richness, abundance or occurrence in the landscape according to the HAH predictions. The values of the response variable are normalized (*S_norm,i_*) so that 100% corresponds to the maximum value found in these landscapes (which happens when the entire local landscape is covered by habitat) and 0% to the case of no habitat in the local landscape. All habitat sites in these examples have however some amount of habitat in their surrounding local landscapes, so that *S_norm,i_* never goes below 15% in these examples. See Appendix S3 for a similar figure but calculating the response variable as a power function of the habitat amount in the local landscape; the numbers of Mean, Max and Q1 vary, but they lead to the same trends and conclusions.

**FIGURE 3.**
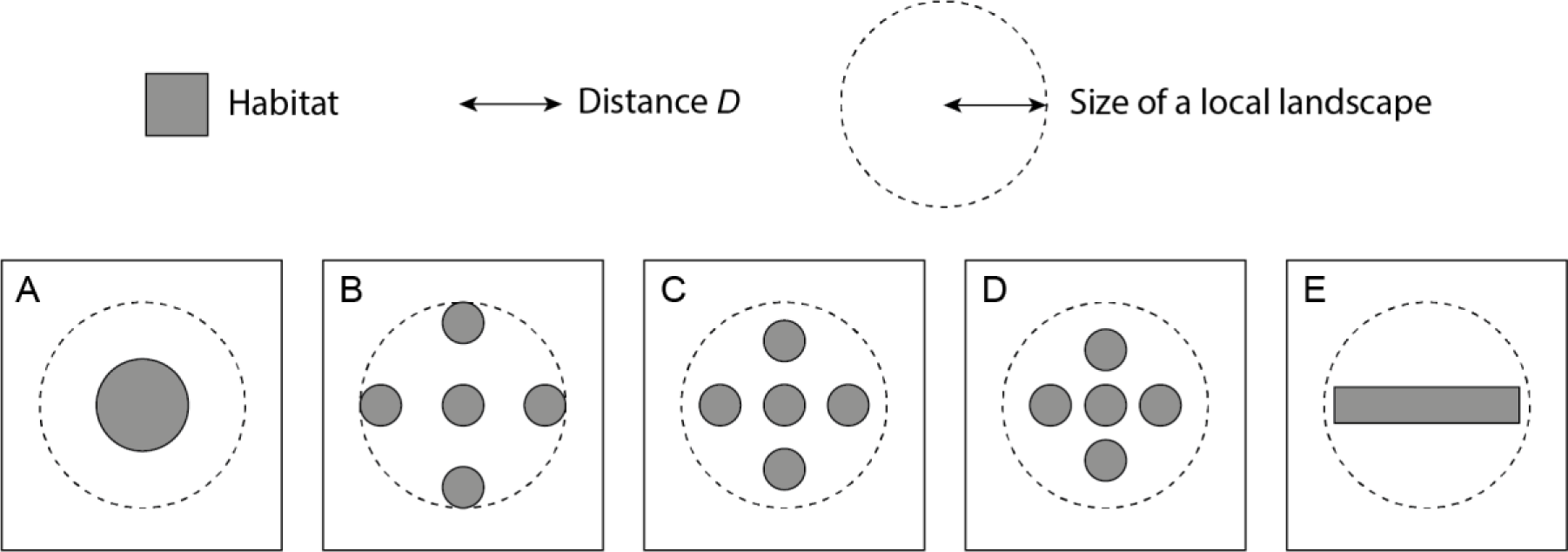
Five landscapes with the same habitat amount but different habitat spatial configuration in which the habitat is all distributed within the extent of a single local landscape of the Habitat Amount Hypothesis, as given by the distance *D* (scale of effect). All habitat in landscapes A and E is found in a single habitat patch, which is more elongated in E. The habitat in landscapes B, C and D is fragmented in five equally sized habitat patches, but the distance between them differs in each of these landscapes. In these examples, it is assumed that no additional habitat is found beyond the boundaries shown for each of the landscapes.

**FIGURE 4.**
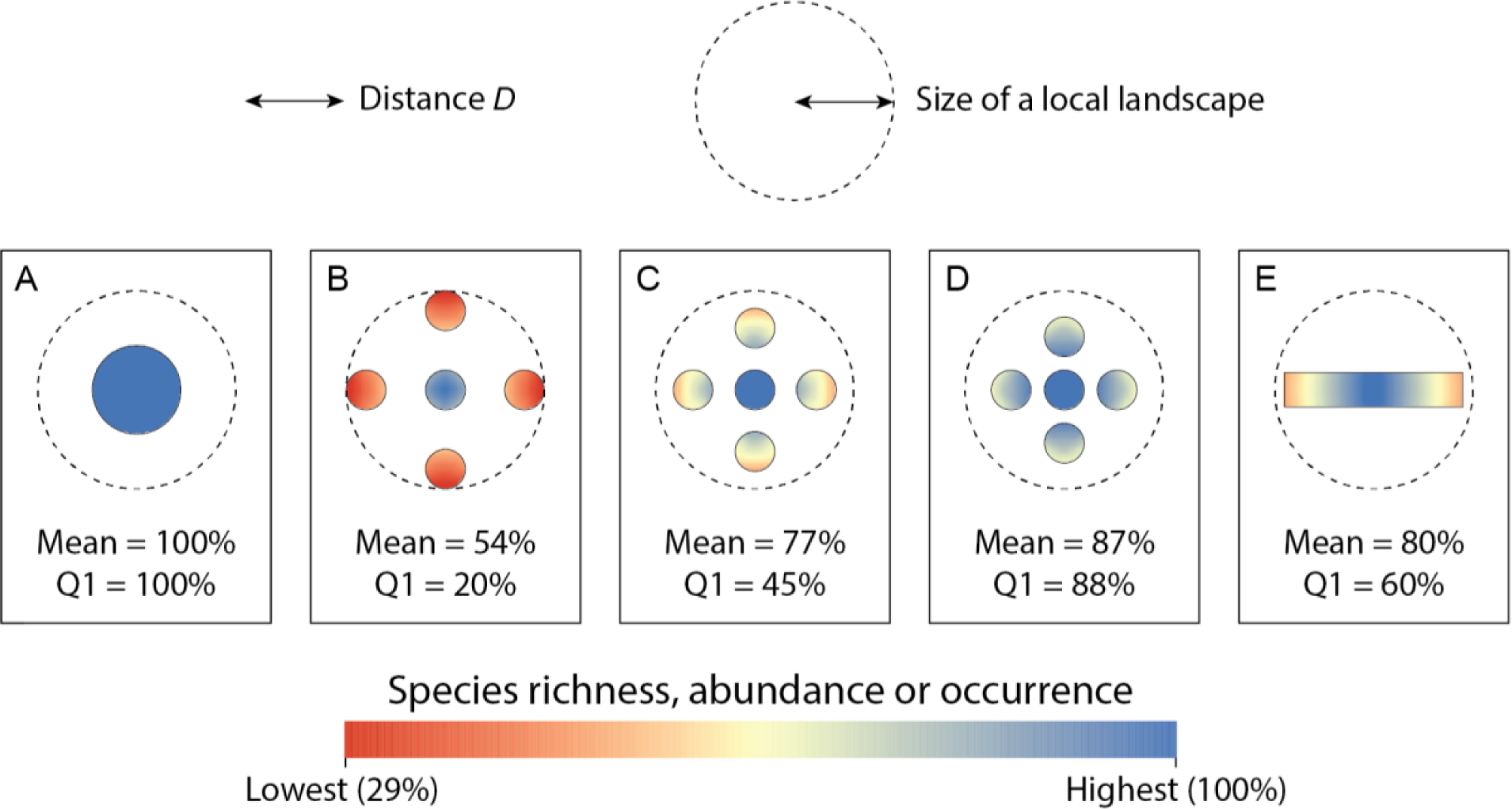
Species richness, abundance or probability of occurrence (*S_norm,i_*) given by the predictions of the Habitat Amount Hypothesis (HAH) in each of the landscapes in Figure 3, i.e. calculated for each location (habitat site) *i* as a linear function of the amount of habitat in the local landscape (distance *D*) around the site, as described in section 2.2. The figure indicates, for each landscape, the mean value (Mean) of *S_norm,i_* in the landscape as predicted by the HAH, as well as the percentage of habitat area in each landscape that has a *S_norm,i_* value within the top quartile (Q1; within the 75-100% range). In all the landscapes there are some habitat sites in which the maximum response variable value is reached (Max = 100%). The results show that an increased habitat fragmentation (breaking apart of habitat), elongation or inter-patch distance, while holding constant the amount of habitat, has negative effects on the species richness, abundance or occurrence in these landscapes according to the HAH predictions. The values of the response variable are normalized (*S_norm,i_*) so that 100% corresponds to the maximum value found in these landscapes (which corresponds to approximately one fifth of the local landscape covered by habitat) and 0% to the case of no habitat in the local landscape. All habitat sites in these examples have however some amount of habitat in their surrounding local landscapes, so that *S_norm,i_* never goes below 29% in these examples. See Appendix S3 for a similar figure but calculating the response variable as a power function of the habitat amount in the local landscape; the numbers of Mean and Q1 vary, but they lead to the same trends and conclusions.

Finally, I calculate the following summary statistics of the response variable values over the entire habitat area in each landscape (aggregated values for all the habitat sites in each landscape): mean (Mean) and maximum (Max) value of *S_norm,I_* and percentage of habitat area in each landscape that has a *S_norm,i_* value within the top quartile (Q1; within the 75-100% range), as shown in Figures 2 and 4.

In summary, I calculate the response variable (species richness, abundance or occurrence probability) in each location (site) in these landscapes using solely the HAH predictions (Figures 2 and 4). That is, I calculate species richness, abundance or occurrence probability in each location (site) exclusively as an increasing function of the amount of habitat in the local landscape around each site. By doing so, I obtain distributions of the response variable (e.g. species richness) that are necessarily in full accordance with the HAH, but examine the broader results for the entire landscape (considering all habitat sites together) rather than narrowly focusing on an individual habitat site only.

All the conclusions obtained in this study hold for other plausible non-linear relationships, such as a power law giving a convex shape of the function between habitat amount in the local landscape and species richness in the site (linear relationship when both variables are log-transformed), as shown in Appendix S3.

### 2.3. The HAH actually implies negative effects of habitat fragmentation on species richness, abundance or occurrence

The predictions of the HAH in these illustrative landscapes (Figures 2 and 4) show clear effects of habitat configuration on species richness, abundance or occurrence (whatever the response variable considered through the HAH). Species distributions, as predicted by the HAH, differ considerably across landscapes with the same total amount of habitat depending on habitat configuration (Figures 2 and 4).

Habitat fragmentation (breaking apart of the habitat into a larger number of smaller patches) in the landscape, while holding constant the amount of habitat, negatively affects, according to the HAH predictions, species richness, abundance or occurrence in all or many of the habitat sites in the landscape (Figures 2 and 4). For instance, the fragmented habitat in Figure 2C (composed of 18 small habitat patches) has an average and maximum value of the response variable of 33% compared to an average of 86% and a maximum of 100% when the same total habitat area is found in a single habitat patch as in Figure 2A. As indicated by the examples highlighted in green in Figure 1, all sites in landscape C have only part of their local landscapes covered by habitat, while in landscape A there are sites with their local landscapes fully covered by habitat. It is possible to decrease the mean and maximum value of the response variable as much as desired by producing habitat configurations that are even more fragmented than that in Figure 2C. This is illustrated in some of the cases in, Figures S4.1 and S4.2 in Appendix S4. For instance, the mean value of the response variable is only 7% in the landscape E in Figure S4.2, in which the habitat patches are five times smaller than those in Figure 2C. While the mean value of the response variable always decreases with more fragmentation in all these cases in Figures 2 and S4.2, the maximum value may or may not be affected depending on the case. For example, the habitat in Figure 2B is more fragmented than in Figure 2A, but there are still some sites (even if less in 2B than in 2A) that attain the same maximum value of the response variable in both landscapes. The percentage of the habitat area with high values of species richness (or abundance or occurrence) decreases however significantly; compare Q1 values in Figures 2A and 2B. Habitat fragmentation also has negative effects on the response variable when considering smaller landscapes with the same extent of a local landscape (Figure 4). However, only the mean value of the response variable and the percentage of habitat area with high values of the response variable show declines at this scale of the local landscape in the cases illustrated in Figure 4. The maximum value of the response variable is not affected by any of the different spatial configurations considered in Figure 4; some of the habitat sites (even if less in the more fragmented patterns) still present the same maximum value of the response variable in these local landscapes (Figure 4).

A more elongated shape of the habitat patches (narrower patches), or larger perforations within the patches, while holding constant the number of patches and the total habitat amount, also imply, according to the HAH predictions, a decline in species richness, abundance or probability of occurrence in all or many of the habitat sites in the landscape. This is apparent by comparing for instance Figure 2H with Figure 2A for the effect of patch elongation, or Figure 2F or 2D with 2A for the effect of patch perforation. These effects are more pronounced when the considered landscapes are larger than the size of a local landscape in the HAH; in several of the cases in Figure 2 it severely affects both the mean and the maximum values of the response variable (Figure 2). They also happen, although to a lower extent and not affecting the maximum values, in landscapes of the same size as a local landscape (Figure 4); compare Figure 4E with Figure 4A.

An increase in the distance between the habitat patches, while holding constant the total habitat amount and the number, size and shape of the patches, can also lead to decreases in species richness, abundance or occurrence probability in the landscape, depending on the cases. In landscapes larger than the scale of effect of the HAH, the inter-patch distance effects can happen but are generally not very strong. This can be seen, for example, in the six habitat patches in Figures 2G and 2B, in which the mean species richness slightly declines from 65% when they are very close to each other to 59% when they are further apart. Any increase in the inter-patch distances beyond those in Figure 2B (all patches are separated by distances larger than *D*), will not produce any additional decline in the response variable. The inter-patch distance effects are more prominent when considered within the extent of a local landscape. For example, the mean species richness in the landscape declines from 87% when the five habitat patches are close to each other (Figure 4D) to 54% when these five patches are separated by a larger distance (Figure 4B).

## 3. WHY HAVE THE IMPLICATIONS OF THE HABITAT AMOUNT HYPOTHESIS BEEN MISINTERPRETED? THE CONFUSION OF DIFFERENT SPATIAL SCALES

### 3.1. The Habitat Amount Hypothesis predicts species richness at the scale of individual habitat sites, which is not the landscape scale

The Habitat Amount Hypothesis (HAH), by definition, implies that species richness (or occurrence or abundance) *in an individual habitat site* is only determined by the amount of habitat *in the local landscape surrounding the site* (distance *D* around the site), regardless of habitat configuration. The HAH, however, *does not imply* that species richness (or occurrence or abundance) in *an entire landscape*, composed of multiple habitat sites with different local landscapes distributed throughout the entire landscape, is only determined by the amount of habitat *in the entire landscape* regardless of habitat configuration. This important distinction has been overlooked in many of the interpretations or applications of the HAH, which has led to incorrect inferences on the implications and reach of the HAH. One of these misinterpretations is, for example, stating that, because the HAH holds, biodiversity protection policies should focus on habitat amount, irrespective of its spatial configuration (Melo et al., 2017). Other similar misinterpretations can be found in the statements on the HAH implications that are summarized in the Introduction. As I have shown above, the HAH does in fact predict, when applied over entire landscapes made up of *multiple habitat sites*, negative effects of habitat fragmentation, elongation or inter-patch distance on species richness, abundance or occurrence in all or many of the habitat sites in the landscape. This holds both when considering landscapes with the same extent as a local landscape (Figure 4) and when considering larger landscapes (Figures 2 and S4.2), as well as when considering patches either smaller or larger than the extent of a local landscape (Figures 2, S4.2 and 4).

### 3.2. A correct and a wrong statement on what the HAH says

It is correct to state that, if the HAH holds, (i) species richness *in an individual habitat site* is affected only by the amount and not by the spatial configuration of habitat *in the local landscape* around that particular site. This statement cannot be however generalized further. It is *not correct* to state that, if the HAH holds, (ii) species richness in *all of the habitat sites* in the landscape is affected only by the amount of habitat in the entire landscape and not by the spatial configuration of habitat *in the entire landscape*.

There is a big difference between statement (i) and statement (ii). Statement (i) is what the HAH says. Statement (ii) is not what the HAH says but what it has been wrongly interpreted to mean or to imply. There is a major change in the scales concerned in each of these two statements. Statement (i) is correct because it refers only to the scales that are directly involved in the HAH as originally proposed, i.e. an individual habitat site and the local landscape around it. Statement (ii) is incorrect because it involves and refers to spatial scales that are outside of the domain of the HAH as originally proposed, i.e. it makes a statement about habitat amount and configuration over an entire landscape, rather than over the extent of an individual local landscape defined specifically for an individual habitat site. The HAH does not make any direct statement on the effects of habitat amount or configuration over an entire landscape; it is constrained to separately consider individual local landscapes around individual habitat sites. Confounding these different scales, and wrongly considering these two different statements as equivalent, is the main reason why the HAH has been significantly misinterpreted to date.

### 3.3. Habitat configuration in the landscape affects the amount of habitat around different habitat sites

I have shown that species richness, abundance or occurrence over the landscape is influenced by the spatial configuration of the habitat over the landscape when the HAH is assumed to hold. This is because the amount of habitat in the local landscapes surrounding each habitat site does depend on the spatial configuration of the habitat, and not just on its amount, at the landscape scale, as shown in Figures 2, S4.2 and 4. In smaller, more elongated or perforated patches, and in some cases also in more distant patches, there will be less habitat sites with a high amount of habitat in their surrounding local landscapes. This makes species richness, abundance or occurrence in these sites decline with habitat fragmentation or with other habitat spatial configuration changes in the landscape (Figures 2 and 4).

When all habitat is found in a single and compact habitat patch, there will be more sites with a high habitat amount in their local landscape. For example, in the single patch in landscape A in Figure 1 there are many sites (those close enough to the patch centre or, in other words, far enough from the patch boundary) that have their local landscape fully covered by habitat, as illustrated in one of the examples in green in Figure 1A. This is no longer possible when patches are smaller than the scale of effect and they are in addition separated by distances larger than *D*, as is the case for instance of landscape C in Figures 1 and 2, or of landscapes B-F in Figures S4.1 and S4.2. Similarly, when the patch is more elongated or perforated (even if with the same total patch area) there will be less or no habitat sites that are far enough from the (inner or outer) patch boundary as to have a large proportion of their local landscape covered by habitat. This is visually clear in the examples of habitat sites and local landscapes in green in Figures 1F or 1H compared to Figure 1A.

There is therefore a direct relationship between the number of habitat sites that are close (closer than *D*) to a patch boundary and the number of sites that will have lower levels of species richness, abundance or occurrence according to the HAH predictions. In other words, the more habitat sites that are located in the inner part of a habitat patch (farther than *D* from the patch boundary), the higher the species response variable will be.

It is important to note that this should not be interpreted as an edge effect in the sense of being relevant only for species that are edge-sensitive and thrive only in the cores of habitat patches. The examples in Figures 1, 3 and S4.1 could equally represent the core habitat once the edge-influenced area was removed, as in figure 8d in Fahrig (2013). The same arguments made here would apply and the same conclusions would hold for the core-dependent species (interior specialists); those core habitat sites that are in the outer part of the core habitat, closer to the boundary with non-habitat (non-core habitat), will have less habitat (core habitat) in their local landscapes, and will hence have lower species richness. In other words, core area specialists will be influenced by the spatial configuration of the core habitat in the landscape, with negative effects of fragmentation and elongation, according to the HAH predictions, in the same way that generalist or edge-affiliated species will be affected by the spatial configuration of their (different) habitat at the landscape scale. The conclusions presented here equally hold whatever the habitat is for a given species or group of species; the same arguments are valid for any habitat that can be identified and mapped as relevant for the species under consideration.

It is relevant to note that the distance *D* (scale of effect) in the HAH can be seen as an explicit recognition of the importance, and of the negative effect of, habitat isolation for species richness, abundance or occurrence. According to the HAH, the distance between habitat sites matters, so that when the habitat sites, or the habitat patches (which contain sites), are located at large distances (compared to *D*) from each other, biodiversity is affected negatively. This is captured in the HAH through the amount of habitat in a local landscape; if sites are farther from each other, there will be less habitat locally, within the extent of the local landscape of each site. This is illustrated by the examples of habitat sites and their local landscapes highlighted in green in landscapes G and B in Figure 1. In the extreme, if all habitat sites are separated by distances larger than *D*, all the habitat that will be found in the local landscape of any of the sites will be just that contained within the individual site. It also follows that if habitat sites or habitat patches are located at any distance larger than *D*, any additional increase in their distance will not translate in any further decline in the species response variable as predicted by the HAH. Beyond *D*, the characteristics (area of habitat) of a given habitat site or patch will have no influence in the characteristics (species richness, abundance or occurrence) of the other habitat site or patch. It is therefore possible to view the distance *D* in the HAH basically as an isolation distance or as a maximum dispersal distance that is conceptually similar to that considered in many connectivity metrics and analyses.

In summary, regarding fragmentation of the habitat in a landscape or region, any discontinuity in the habitat compared to the case of a single compact habitat patch with the same total habitat area will translate into negative effects on species richness, abundance or occurrence (see Figures 2, 4 and S4.2). This is because the discontinuity will cause an increase in the distance that separates some of the habitat sites, which means that some of these sites will have less habitat in their local landscapes (as they will contain the non-habitat discontinuity rather than the continuous expanse of habitat). Any increase in the discontinuities in the habitat distribution (i.e. any increase in the number of patches for the same total habitat area) will have negative effects on the response variable, up to the point of the distance *D* of the scale of effect. Any further increase in the non-habitat distance or discontinuity between the habitat sites beyond *D* will not translate into further declines in the response variable, as noted above. The HAH implies that the lowest values of species richness, abundance or occurrence in a landscape are obtained, for a given amount of habitat, when all habitat sites are isolated from each other, i.e. when all habitat patches are individual habitat sites located at a distance larger than *D* from each other (so that there is no other habitat site within the local landscape of each site). The opposite configuration extreme is, obviously, when all habitat sites are found in a single compact (circular) patch with the same total habitat area.

### 3.4. The Habitat Amount Hypothesis predicts species richness at a different scale from that at which habitat amount is quantified

It is important to remark that the HAH predicts species richness at a different scale from that at which habitat amount is quantified. The HAH only deals with, and only gives direct predictions on, species richness, abundance or occurrence at the scale of the habitat site. The predictions of the response variable (species richness, abundance or occurrence in a habitat site) are obtained from a single explanatory variable, the habitat amount in the local landscape. The extent or scale of effect involved in the explanatory variable (local landscape for the distance *D*) is different and much larger than the extent of the habitat site involved in the response variable. This difference in the extent of the site (scale of the response variable) and of the local landscape (scale of the explanatory variable) is clear and explicit in the description of the HAH (Fahrig, 2013). However, the meaning and implications of this scale difference for the interpretation and use of the HAH have not been sufficiently emphasized, and in some case have been overlooked, in the literature on the HAH. Fundamentally, this difference in the scale of the species richness and habitat amount variables in the HAH means that the HAH cannot be used to make any direct statement on species richness at any scale different from (larger than) the site scale, including the scale of the local landscape. This condition has been often not kept in mind in the uses, applications and tests of the HAH, which has led to other significant and unreported misinterpretations of the HAH that are described next in Sections 4 and 5.

## 4. THE HABITAT AMOUNT HYPOTHESIS IS COMPATIBLE WITH HIGHER SPECIES RICHNESS OR ABUNDANCE FOR A SINGLE LARGE PATCH THAN FOR SEVERAL SMALL HABITAT PATCHES (SLOSS)

One of the long-standing debates in ecology is the Single Large Or Several Small (SLOSS) debate (e.g. Wilcox & Murphy, 1985). To maximize species richness, or species abundance or occurrence, is it better to have a single large habitat patch or several small habitat patches with the same total habitat area?

Fahrig (2013) stated, in the legend of figure 5 on the empirical evaluation of SLOSS in her paper, that the HAH “predicts that species number should increase with total area, irrespective of the number of patches making up that total”, i.e. that, according to the HAH, the same number of species should be found in a large patch as in several small patches with the same total area. Therefore, Fahrig (2013) considered the HAH to be ‘SLOSS neutral’, which is the term I will use here to refer to the case in which a large patch has the same species richness as several small patches with the same total area.

**FIGURE 5.**
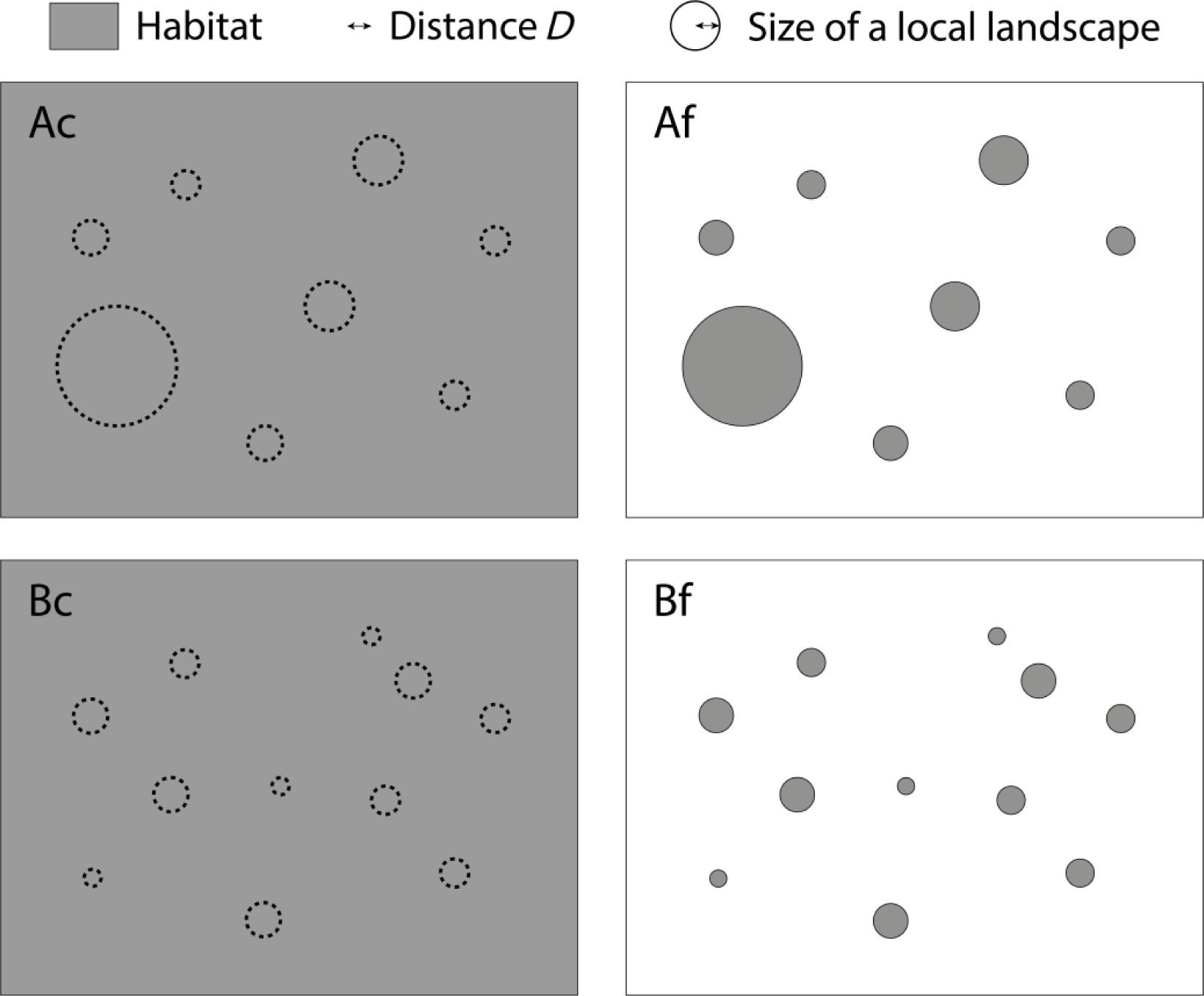
Two sets of different-sized non-continuous patches in fragmented landscapes (Af and Bf) compared to a set of sample areas (dashed lines) equal in size to these patches but contained within a landscape with continuous habitat (Ac and Bc). In case A, some of the habitat patches (Af) and sample areas (Ac) are larger than the size of a local landscape of the Habitat Amount Hypothesis (HAH), as given by the distance *D* (scale of effect). In case B, however, all habitat patches (Bf) and sample areas (Bc) are smaller than the size of a local landscape of the HAH.

The statement of Fahrig (2013) on the HAH implying SLOSS neutrality does not always hold however according to the predictions of the HAH itself. Consider for instance the single large patch in Figure 2A and compare it with the 18 smaller patches with the same total area in Figure 2C. According to the HAH predictions applied to all sites in these landscapes, Figure 2C has a much lower mean and maximum species richness than Figure 2A. Figure 2C has no habitat site with a species richness in the top quartile (75-100%), compared to about two thirds of the habitat area in the top quartile of species richness in the large patch in Figure 2A. Total species richness in the landscape would also be expected to decline in Figure 2C compared to Figure 2A, although the particular values will depend on species turnover, as discussed in Appendix S5.

The same lack of SLOSS neutrality happens for the abundance or occurrence of the individual species to which the HAH applies. If the abundance and/or occurrence of each of the individual species is considerably lower in several small patches than in a large one, as is apparent in the comparison between Figures 2A and 2C, it can only happen that less of these species are found in the small patches (all together) than in the single large patch. In other cases, such as in the comparison of Figures 2A and 2B, or of Figures 4A and 4B, the mean values of the response variable decreases considerably in the small patches compared to the large patch, but there are still a few habitat sites (20% or less) in the fragmented habitat that have the same maximum level of the response variable as in most of the habitat sites in the single large patch to which they are compared. In these cases, the total species abundance and occurrence in the landscape will decline, but the total species richness in the landscape may be the same in both of the SLOSS situations, given that such maximum species richness is found in some of the habitat sites in all these cases. See a more detailed discussion on this in Appendix S5.

Taken together, these cases show that the HAH does not imply, in general, that several small patches will have the same species richness, abundance or occurrence probability as a large patch with the same total area. There are many cases in which these response variables will be, according to the HAH predictions, higher in a single large habitat patch than in several small ones with the same total area. Fahrig (2013) considered that having more species in a single large patch than in several small patches of the same total area would be indicative of an island effect consistent with the island biogeography theory (IBT). I here show, however, that the HAH is compatible with that same result and, therefore, whether the HAH or the IBT hold cannot be judged simply based on this SLOSS comparison without any additional, more detailed consideration. If, in general, the same number of species is found in a set of fragmented patches than in a single large patch of the same total area, the explanation has to be found in a theory or hypothesis different from the HAH.

## 5. A STEEPER SLOPE OF THE SPECIES-AREA RELATIONSHIP FOR MORE FRAGMENTED HABITAT IS COMPATIBLE WITH THE HABITAT AMOUNT HYPOTHESIS

Species-Area Relationships (SARs), which describe how the number of species recorded increases with the sampled area, are one of the most important laws in ecology (Scheiner, 2003; Triantis, Guilhaumon, & Whittaker, 2012).

Fahrig (2013) stated that the HAH can be tested by comparing the slope of the SAR across a set of different-sized patches with the slope of the SAR across a set of sample areas equal in size to these patches but contained within a region of continuous habitat. A steeper slope for the fragmented patches would be, according to Fahrig (2013), consistent with the island effect of the island biogeography theory (IBT) and inconsistent with the HAH. Other studies have followed this way of testing the HAH proposed by Fahrig. These studies either concluded support for the HAH because of no difference in the SAR slope in that comparison (Rabelo et al., 2017) or rejected the HAH because they found a steeper slope of the SAR for the non-continuous patches (Haddad et al., 2017; Bueno & Peres, 2019).

I here show, however, that the HAH predictions lead to species richness patterns that differ considerably between habitat patches and samples areas with the same size in continuous habitat. This is clearly illustrated in the landscapes in Figure 5 and in the results of Figure 6, which I obtained using the same calculation procedure as described in section 2.2 (i.e., the same procedure applied above to produce Figures 2, 4 and S4.2). This difference in the predictions can be such that both mean and maximum species richness decrease in the fragmented habitat patches compared to the equivalent sample areas in continuous habitat (compare Figures 6Bf and 6Bc). Alternatively, it can be such that mean species richness decreases but there are some (even if much less) habitat sites with the same maximum value of species richness as in the non-fragmented case (compare Figures 6Af and 6Ac).

**FIGURE 6.**
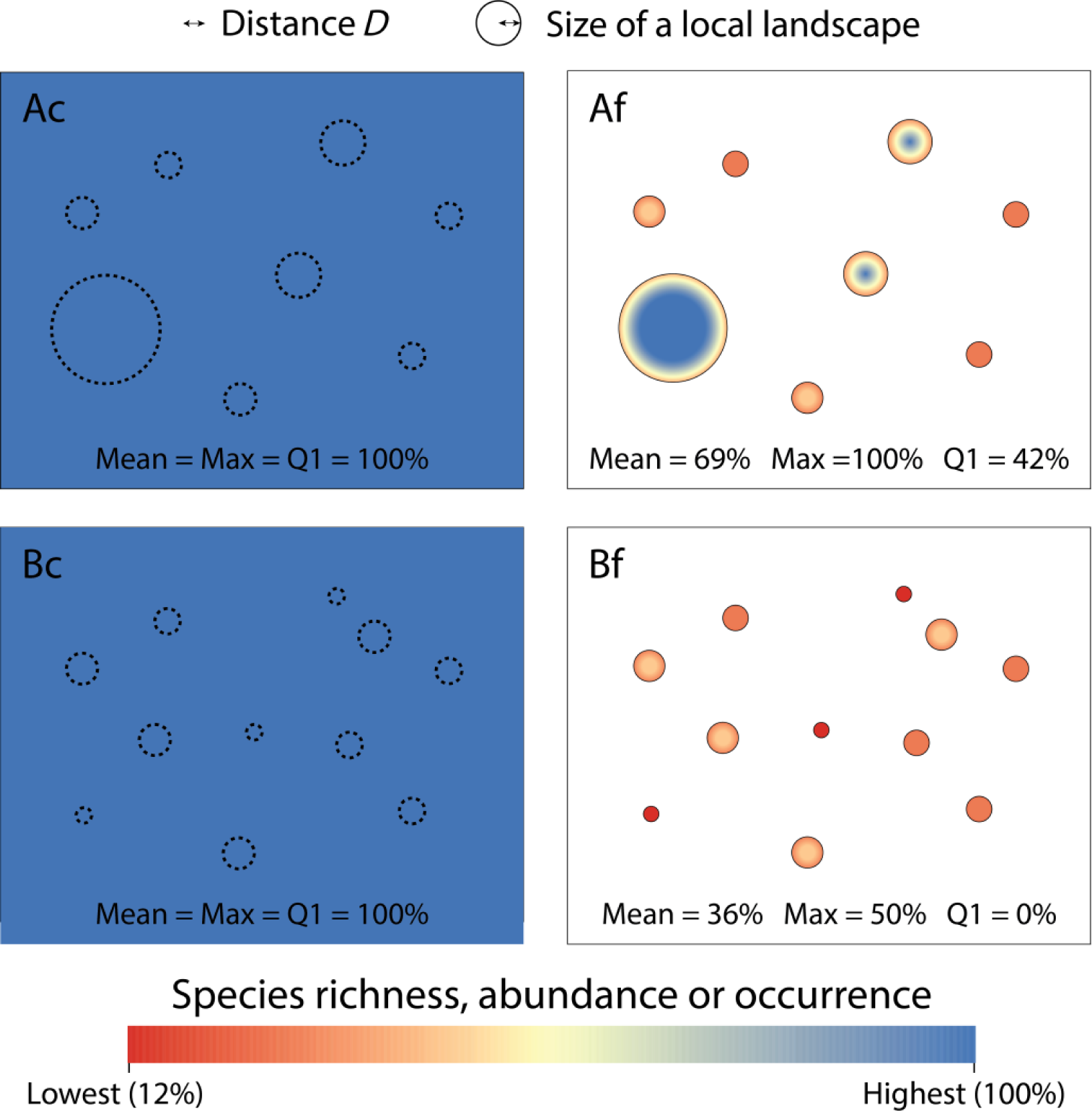
Species richness, abundance or probability of occurrence (*S_norm,i_*) given by the predictions of the Habitat Amount Hypothesis (HAH) in each of the landscapes in Figure 5, i.e. calculated for each location (habitat site) *i* as a linear function of the amount of habitat in the local landscape (distance *D*) around the site, as described in section 2.2. The figure indicates, for each landscape, the mean (Mean) and maximum (Max) value of *S_norm,i_* in the landscape as predicted by the HAH, as well as the percentage of habitat area in each landscape that has a *S_norm,i_* value within the top quartile (Q1; within the 75-100% range). In the landscapes with continuous habitat (Ac and Bc), the habitat is assumed to extend beyond the boundaries of the shown rectangles, and hence the response variable remains high even close to the rectangle border. The values of the response variable are normalized (*S_norm,i_*) so that 100% corresponds to the maximum value found in these landscapes (which happens when the entire local landscape is covered by habitat) and 0% to the case of no habitat in the local landscape. All habitat sites in these examples have however some amount of habitat in their surrounding local landscapes, so that *S_norm,i_* never goes below 12% in these examples. See Appendix S3 for a similar figure but calculating the response variable as a power function of the habitat amount in the local landscape; the numbers of Mean, Max and Q1 vary, but they lead to the same trends and conclusions.

Given the difference in species richness in the different-sized patches in Figure 6Af and 6Bf, and the lack of any difference in species richness in the comparable sample areas in continuous habitat (Figure 6Ac and 6Bc), all of which result from the HAH predictions, it follows that the SAR slope for the fragmented patches has to be steeper than for the continuous habitat. Indeed, in the habitat patches there is an increase in species richness as larger areas (larger patches) are considered, while this is not the case in the equivalent sample in continuous habitat, according to the HAH predictions (Figure 6). This applies to a SAR of Type IV (Scheiner, 2003), in which each data point is from a sample corresponding to a unique area (patch), but would also hold for other SAR types, such as those constructed by estimating the mean diversity for a given area from multiple combinations of habitat sites (Scheiner, 2003). See Appendix S5 for some more detailed considerations in this regard.

Therefore, the HAH is compatible with, and implies in many cases such as those illustrated in Figure 6, different SAR slopes depending on the degree of habitat fragmentation, and steeper SAR slopes for different-sized patches than for sample areas equal in size to these patches but contained within continuous habitat. This highlights the need to revisit the ways in which the HAH can be tested, as well as the support, or the lack of support, to the HAH that has been concluded in previous studies based on the slope of the SAR.

For instance, Bueno and Peres (2019) found that the slope of the SAR was steeper for islands surrounded by low habitat amounts than for islands surrounded by high habitat amount. Based on this finding, and on Fahrig (2013), they concluded that the IBT, and not the HAH, was supported in their case. Similarly, Haddad et al. (2017) found that the slope of the SAR was steeper in fragmented than in more continuous landscapes, and concluded that this result provided a direct refutation of the HAH. However, I have here shown that the HAH predicts species richness patterns that lead to steeper slopes when the habitat patches are isolated (surrounded by non-habitat areas) than when they are surrounded by continuous habitat (Figure 6). Therefore, this specific result by Haddad et al. (2017) or Bueno and Peres (2019) cannot be used to support IBT against HAH. This specific result does indeed highlight the importance of habitat configuration and fragmentation for species richness, but paradoxically is not in contradiction with the HAH, but is in agreement with a prediction that arises from the HAH itself.

Conversely, Rabelo et al. (2017) found similar slopes in the SAR for fragmented patches than for comparable sample areas in continuous habitat. They interpreted this result as providing stronger support for the HAH than for the IBT, which was concluded not to explain species richness in their study system. However, given that the HAH is compatible with steeper SAR slopes in fragmented than in continuous habitat, as illustrated from the results in Figure 6, this finding from Rabelo et al. (2017) cannot be used to support HAH against the IBT.

The other ways to test the HAH proposed by Fahrig (2013) different from the SAR slope, which are summarized in figures 7 and 11 in her paper, remain appropriate for testing the HAH. This is so because, unlike for the SAR slope, these other ways of testing the HAH refer exclusively to predictions at the habitat site level that are made from the amount of habitat in the local landscape surrounding the site, which are respectively the response and explanatory scales involved directly in the HAH predictions.

## 6. HANSKI’S RESPONSE TO FAHRIG

Hanski (2015) elaborated, in his reply to Fahrig (2013), on four objections to the HAH, the first of which was: “My first concern is that the spatial scale of the habitat amount hypothesis is the local landscape around an individual study plot. This is a narrow perspective, which does not allow one to address fundamental questions about the occurrence of species within large landscapes with more or less habitat that is more or less fragmented”. This is indeed an important point, to which I would only add two notes. First, that there are two spatial scales, and not just one, in the HAH: the scale of the local landscape, where the habitat amount is measured, and the scale of the habitat site (study plot as named by Hanski) in which species occurrence, abundance or richness is predicted. Second, that even when the HAH only predicts the occurrence of species in individual (small) habitat sites, it is possible to use the HAH, if one assumes that it holds, to predict species occurrence in each of the sites over large landscapes, providing the HAH-based pattern of species occurrence within large landscapes with more or less fragmented habitat, as done here in Figures 2, 4 and 6.

Hanski (2015) continued, stating that the conclusion on the unimportance of habitat configuration related to the HAH “may be valid for small spatial scales and when the total amount of habitat is large, but modelling and empirical studies demonstrate adverse demographic consequences of fragmentation when there is little habitat across large areas”. He continued with the following example: “to see the significance of this distinction, consider a focal site with its local landscape, which in one case is completely isolated from the rest of the habitat in the landscape, in another case completely surrounded by other habitat. It would be most surprising if the landscape context were not to make a difference to the occurrence of the species”. Hanski (2015) did not elaborate further on this issue. Particularly he did not notice, and this has not been noted either in any other article on the HAH so far, that the HAH itself, regardless of any other modelling approaches or empirical studies, does predict the importance of habitat configuration (or landscape context) in cases such as the one described by Hanski (2015). In fact, the example given by Hanski (2015) is demonstrated as true even if assuming that the HAH holds and using the species occurrence predictions provided by the HAH: compare the patches in Figures 6Af and 6Bf, which are ‘isolated’ from (not adjacent to) the rest of the habitat in the landscape, with the sample areas of the same size in Figures 6Ac and 6Bc, respectively. The landscape context does make a difference, and the isolated patches show much lower levels of species occurrence or abundance than their equivalent sample areas surrounded by habitat (Figure 6). This happens in the particular case of a patch with the same size as a local landscape (Figure 6A), which was the one specifically mentioned by Hanski in his example, but also for other patches larger and particularly smaller than a local landscape (Figure 6). Landscape context and habitat spatial configuration do make a difference, and have effects that are distinct from and go beyond those of habitat amount. This conclusion is reached without needing an alternative hypothesis to the HAH or other mechanisms or theories such as island biogeography or metapopulation theory; this is the prediction that arises from the Habitat Amount Hypothesis itself.

## 7. CONCLUSIONS

The Habitat Amount Hypothesis (HAH) has triggered an intense debate, and a considerable number of empirical studies, on the relative importance of habitat amount and habitat fragmentation for biodiversity patterns and persistence. The prevailing view and interpretation of the HAH has been that, if or where it holds, it negates the existence of fragmentation effects and implies that conservation strategies should focus on retaining the maximum overall amount of habitat regardless of its configuration (Fahrig, 2013; Haddad et al., 2017; Melo et al., 2017; Torrenta & Villard, 2017; MacDonald et al., 2018; Bueno & Peres, 2019), as summarized in the Introduction.

I have here shown that, contrary to the current interpretations of the HAH, the predictions of the HAH actually imply clearly negative effects of habitat fragmentation on species richness, abundance or occurrence in a landscape or region, and that these fragmentation effects are distinct from those of habitat amount in the landscape or region. I have shown that increased habitat fragmentation, elongation, perforation or (in some cases) inter-patch distance, while holding constant the amount of habitat, have negative effects on species richness, abundance or occurrence in all or many of the habitat sites in the landscape according to the HAH predictions. Therefore, even if the HAH holds, conservation strategies cannot focus only on the amount of habitat in a landscape or region, and should also consider its spatial configuration.

Many of the misunderstandings of the HAH arise from the omission of the key difference between the spatial scales involved in the HAH and the landscape or regional scales at which conservation management actions are to be applied. The HAH is based on measuring the amount of habitat in the local landscape (scale of effect) around a given habitat site, but only provides direct predictions on species richness or occurrence at the scale of the individual habitat sites, which are much smaller than the landscape scale, the local landscape scale or the patch scale. When the site-scale HAH predictions are applied over entire landscapes, rather than to individual habitat sites considered separately, the important and distinct effects of habitat configuration clearly show up. The difference in the scale of the species richness and habitat amount variables in the HAH means that the HAH cannot be used to make any direct statement on species richness at any scale different from (larger than) the site scale, including the scale of the local landscape. This condition has often not been kept in mind in the use, application and conceptualization of the HAH, which has led to significant and previously unreported misinterpretations of the implications of this hypothesis and of the ways it can be actually tested.

I reiterate that I have here not provided any argument in favour or against the validity of the HAH or its underlying ideas. My arguments only refute some of the interpretations that have been made regarding the meaning or implications of this hypothesis, but not the HAH itself, which I have assumed to hold in all my argumentations. My conclusions do not mean that concepts behind the HAH logic, such as considering equally-sized habitat sites instead of habitat patches as the natural spatial units for measuring and analysing species richness (Fahrig, 2013), are not of value for landscape ecology and biogeography. Even some empirical studies that have not found support for the HAH have agreed with the interest of shifting from habitat patches to equally-sized units of analysis (Torrenta & Vilard, 2017).

Hanski (2015) concluded, in his response to Fahrig (2013) objecting the validity of the HAH, that “habitat fragmentation poses a threat to biodiversity, in addition to the threat posed by the loss of the total amount of habitat. Fragmentation effects should not be overlooked in ecology or in conservation”. Haddad et al. (2017) concluded, in their study rejecting the HAH, that “in conservation, only considering one attribute of landscapes (i.e. habitat amount) … could misguide protected area management”. I support these conclusions, and show that they are, in fact, a prediction of the HAH. Reaffirming the importance of habitat fragmentation and configuration for biodiversity does not depend on rejecting the HAH and using other explanatory framework, hypothesis or theory; on the contrary, the actual implications and predictions of the HAH are a clear support for the importance of habitat fragmentation and configuration for biodiversity.

## ACKNOWLEDGEMENTS

I thank three colleagues for their helpful comments on an earlier version of this manuscript and Laurent Bergès for stimulating me to think in more detail about the Habitat Amount Hypothesis by inviting me to give the opening talk of the landscape ecology symposium he organized in the International Conference on Ecological Sciences (Sfecologie) held in Rennes (France) in October 2018.

## APPENDIX S1. Results of the analyses for different sizes of the habitat sites

In all the examples and figures shown in the main text (Figures 1-6), the habitat distribution has been represented through a raster layer in which habitat sites correspond to equally-sized squared cells with a size of 0.005*D* x 0.005*D*, where *D* is the radius of a circle around the center of the cell (habitat site) that defines the local landscape (scale of effect) for each habitat site. The predictions of the HAH were applied to these habitat sites (squared cells) as described in section 2.2 in the main text. This cell size (habitat site size) was small enough as to avoid any visible ‘pixelation’ effects and to provide a smooth representation of the habitat when rasterizing the different circular and rectangular patch shapes in the landscapes in Figures 1, 3 and 5, and of the resultant species distribution patterns in Figures 2, 4 and 6.

All the results and conclusions obtained, and the arguments made using the examples in Figures 1-6 equally hold if other sizes of the habitat sites (cell sizes) are considered, with only minor differences in some of the detailed numbers, which are inconsequential for the purposes of this study. This is shown clearly by considering, for example, habitat sites with a size of 0.05*D* x 0.05*D*, i.e. sites with an area that is 100 times larger than that considered in Figures 1-6. The results of considering this habitat site size of 0.05*D* x 0.05*D* are shown in Figures S1.1, S1.2 and S1.3, which can be compared to those of Figures 2, 4 and 6 in the main text (habitat site size of 0.005*D* x 0.005*D*), respectively. Only minor or very minor differences can be found in this comparison. In comparing Figure S1.1 with Figure 2, hardly any difference can be reported. In Figure S1.2, a more pixelated pattern is apparent compared to the smoother results in Figure 4 in the main text, but there are no noticeable variations in the summary values (Mean, Q1) between the two figures. Similar comments apply to the comparison of Figures S1.3 and 6.

**FIGURE S1.1.**
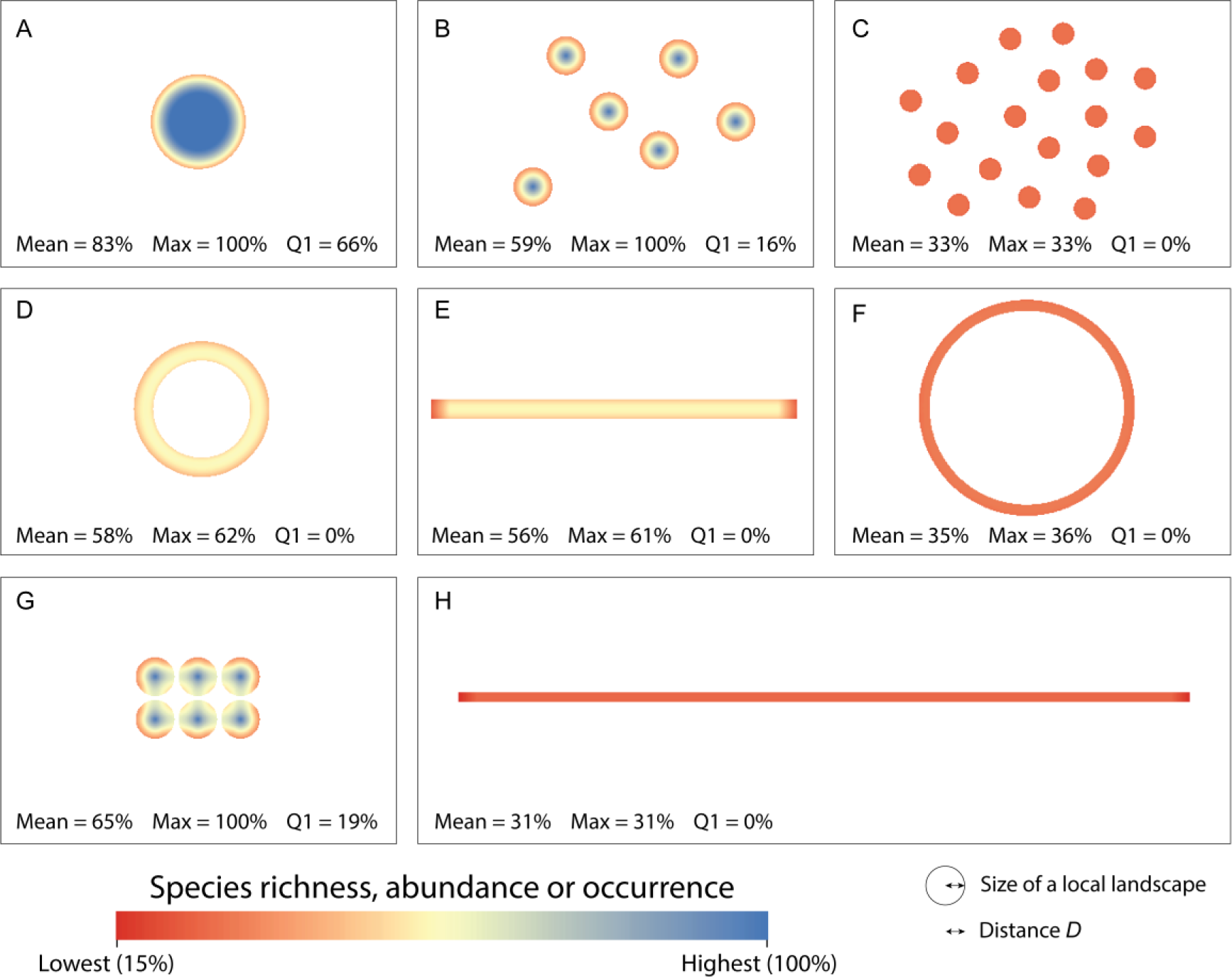
Species richness, abundance or probability of occurrence (*S_norm,i_*) given by the predictions of the Habitat Amount Hypothesis (HAH) in each of the landscapes in Figure 1 when the size of the habitat sites (squared habitat cells) is 0.05*D* x 0.05*D* (compared to habitat sites of size 0.005*D* x 0.005*D* in Figure 2 in the main text). All other specifications are as in Figure 2 in the main text, including the use of a linear relationship between *S_norm,i_* and the amount of habitat in the local landscape around site *i*.

**FIGURE S1.2.**
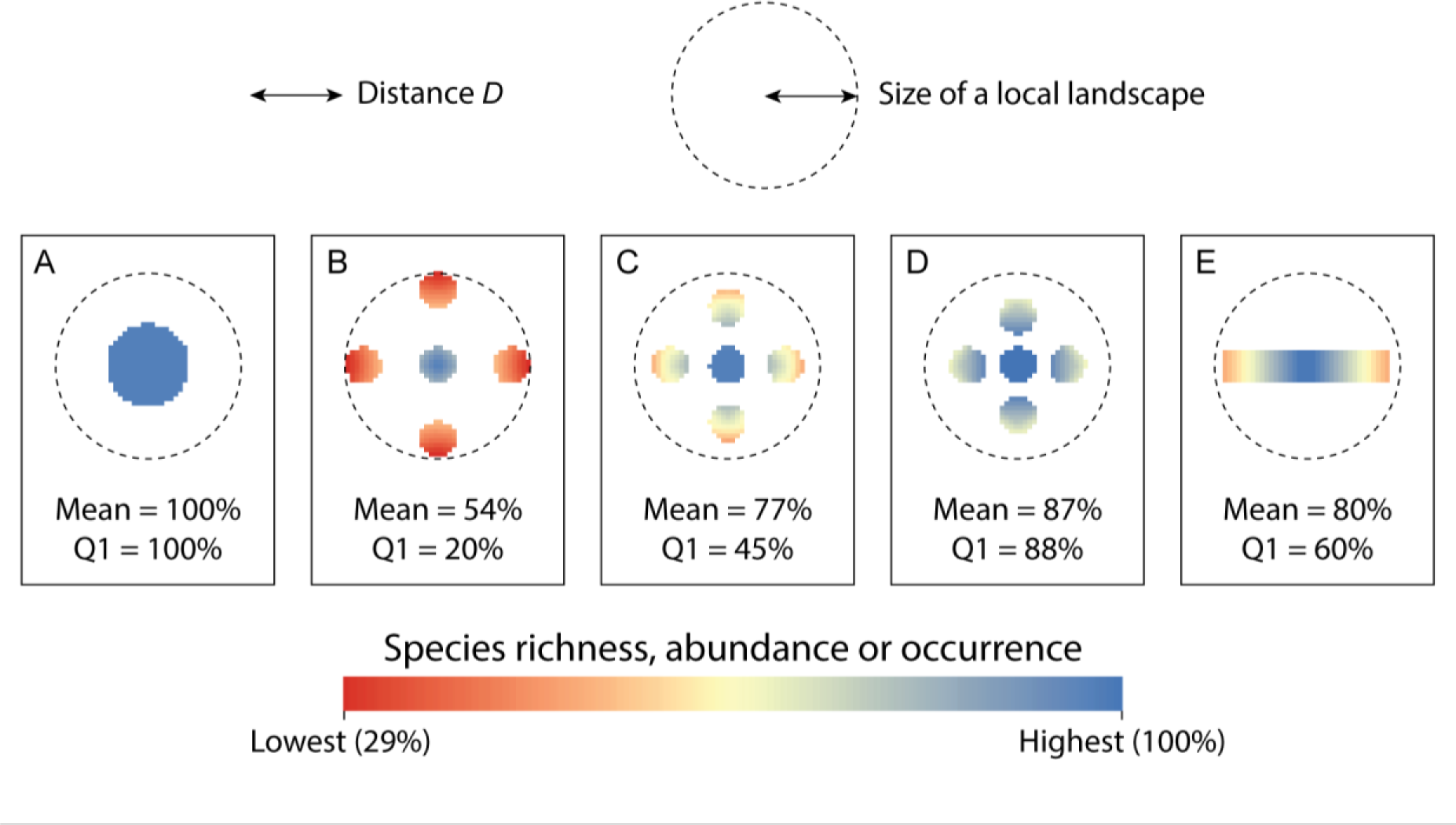
Species richness, abundance or probability of occurrence (*S_norm,i_*) given by the predictions of the Habitat Amount Hypothesis (HAH) in each of the landscapes in Figure 3 when the size of the habitat sites (squared habitat cells) is 0.05*D* x 0.05*D* (compared to habitat sites of size 0.005*D* x 0.005*D* in Figure 4 in the main text). All other specifications are as in Figure 4 in the main text, including the use of a linear relationship between *S_norm,i_* and the amount of habitat in the local landscape around site *i*.

**FIGURE S1.3.**
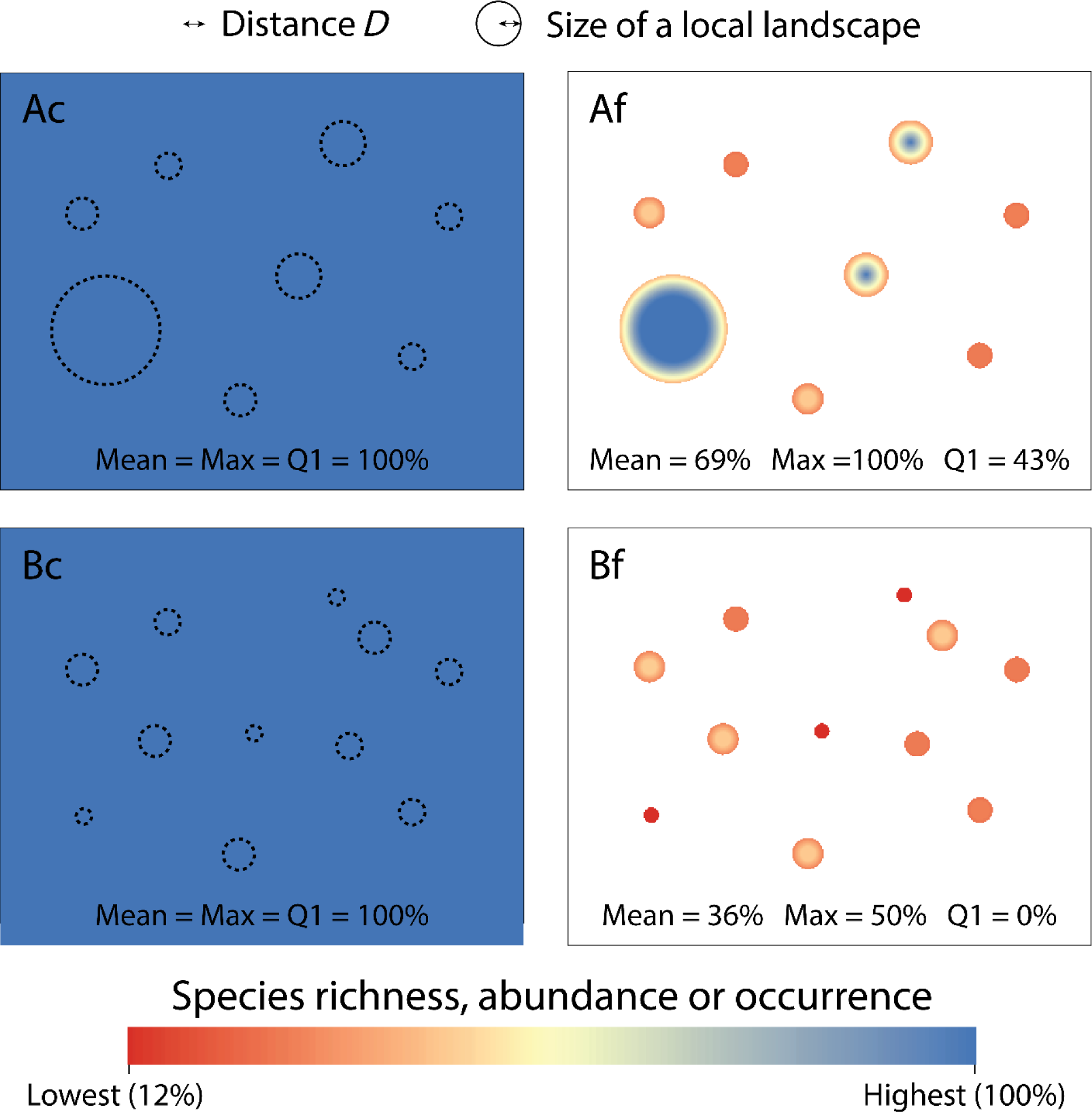
Species richness, abundance or probability of occurrence (*S_norm,i_*) given by the predictions of the Habitat Amount Hypothesis (HAH) in each of the landscapes in Figure 5 when the size of the habitat sites (squared habitat cells) is 0.05*D* x 0.05*D* (compared to habitat sites of size 0.005*D* x 0.005*D* in Figure 6 in the main text). All other specifications are as in Figure 6 in the main text, including the use of a linear relationship between *S_norm,i_* and the amount of habitat in the local landscape around site *i*.

## APPENDIX S2. The size of the habitat patches and the Habitat Amount Hypothesis

Fahrig (2013) did not specify any limitation, nor imposed any restriction, on the size of the habitat patches that could be considered or present in a given study area so that the HAH would remain valid and applicable. It is true that in most of the schematic figures in Fahrig (2013), the drawn habitat patches are smaller than the scale of effect or local landscape (circle with radius *D*). These are however illustrative examples with simple schematic settings designed for clarity, which may not be interpreted as implying a general statement or formal formulation in this regard. For instance, in the illustrative figure 9 in Fahrig (2013), the examples of circles of radius *D* (drawn as potential candidates for the scale of effect) include circles that are either smaller or larger than some of the habitat patches. In fact, most of the studies on the HAH in real-world landscapes have considered, when intending to determine the value of *D* (distance with the strongest correlation between habitat amount and the response variable), ranges of candidate distances *D* that included values smaller than some of the patches in the analysed landscapes (Melo et al., 2017; Rabelo et al., 2017; Torrenta & Villard, 2017; Gardiner et al., 2018; Vieira et al., 2018; Bueno & Peres, 2019). From these studies, the finally selected *D* was smaller than some of the habitat patches in Torrenta and Villard (2017), Gardiner et al. (2018) and Bueno and Peres (2019), while the opposite was found in Melo et al. (2017) and Rabelo et al. (2017). This shows that, in general, the HAH has not been interpreted and used as purposefully or necessarily limited to cases where all habitat patches are smaller than the extent of a local landscape (scale of effect *D*). The contrary would considerably detract from the potential usefulness of the HAH, by prescribing it as not applicable to a large number of real-world landscapes. In addition, the logic behind the HAH is to move away from a patch-based analysis of the landscape, and to use equally-sized habitat sites rather than patches as the spatial units for measuring and analysing species richness (Fahrig, 2013). It would be therefore in contradiction with this logic to impose a patch-based restriction (a patch-area based restriction) to the domain of applicability of the HAH, which is proposed as independent from the concept of patches and states that the size of the habitat patch in which a particular site is located has no distinct effect on the species richness of the site. Analogously, the landscapes or regions over which the HAH may be applied are not constrained to be smaller than the scale of effect. Since the habitat patches can be (either individually or in combination) larger than the scale of effect, the same is true for the landscapes or regions in which those patches are found.

## APPENDIX S3. The response variable calculated using a power law: results for the illustrative landscapes S3.1. The function relating habitat amount to species richness in the HAH: is it linear or not?

The formulation of the Habitat Amount Hypothesis (HAH) does not explicitly indicate, nor the literature on the HAH specifically demonstrates, which is the best functional form of the relationship (e.g. linear or non-linear) between the predictive variable (habitat amount in the local landscape) and the response variable (species richness, abundance or occurrence in the habitat site). In the literature on Species-Area Relationships (SARs), the power law is the most widely used and supported (Scheiner, 2003; Triantis et al., 2012), which may suggest its use also in the context of the HAH. However, it is important to note that the HAH is a different case from the SARs, for two main reasons.

First, in SARs, both the response variable (species richness) and the explanatory variable (sampled area) are measured over the same scale or extent. On the contrary, the HAH involves relating two different scales: the scale at which species richness is measured (habitat site) is different and much smaller than the scale at which the area of habitat is measured (local landscape around the site).

Second, the SAR studies typically use sample areas that extend over several orders of magnitude. The range of sampled areas (difference between the maximum and minimum sampled areas) has a mean value of nearly 100,000 km^2^ (91,941 km^2^) and a median of 733 km^2^ in the 601 datasets from 312 SAR studies reviewed by Triantis et al. (2012); see appendix S2 in that paper. About two thirds (68%) of these SAR datasets considered ranges of sampled areas larger than 100 km^2^, and almost half of them (45%) considered ranges of sampled areas larger than 1,000 km^2^; see appendix S2 in Triantis et al. (2012). This is very different from the range of areas considered in the HAH studies, where the habitat area is much smaller and is constrained by the size of the local landscape. In the HAH studies, the local landscapes have a radius *D* ranging from 40 m to 5500 m (Evju & Sverdrup-Thygeson, 2016; Lindgren & Cousins, 2017; Melo et al., 2017; Rabelo et al., 2017; Seibold et al., 2017; Torrenta & Villard, 2017; Thiele, Kellner, Buchholz, & Schirmel, 2018; Vieira, Almeida-Gomes, Delciellos, Cerqueira, & Crouzeilles, 2018; Bueno & Peres, 2019), with an average value of *D* of about 1000 m, which corresponds to an area of about 3 km^2^. Therefore, the habitat areas considered in the HAH literature generally extend over much smaller area ranges than in the SAR studies. This difference has implications, because Triantis et al. (2012) reported that in the SAR studies that considered areas that extended over smaller orders of magnitude, the SARs were best described by a linear relationship, rather than by a non-linear (convex or sigmoid) functional form; see figure 3 in Triantis et al. (2012). A linear SAR appears more characteristic and the best shape for cases spanning a small range of area values (Triantis et al., 2012). This may be an argument in favour of using, in the context of the HAH, a linear (and most parsimonious) function between the response variable (which may be species richness, but also species abundance or occurrence) and the habitat amount in the local landscape, as done in the main text.

In any case, I here show that the conclusions in the main text are general and not affected by the specific functional form of the plausible relationship between the habitat amount in the local landscapes and the response variable in the habitat sites (as long as it is a monotonically increasing function). For this purpose, I here present the results of applying the same calculations for the illustrative landscapes in Figures 1, 3 and 5 in the main text, but now using a power function (instead of the linear one used in Figures 2, 4 and 6 in the main text).

### S3.2. The power law and its exponent (slope parameter)

The power function has the following form relating the response variable (*S_i_*), such as species richness, in habitat site *i* with the habitat amount (habitat area) in the local landscape around *i* (*A_i_*):

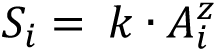

This power function is frequently made linear through a logarithmic transformation of both *S_i_* and *A_i_*, giving:

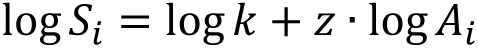

Where *z* is the exponent of the power function, most often referred to as the slope of the log-transformed equation, and *k* is a constant that determines the intercept of the log-log equation.

As for the calculations in the main text, I normalize the values of the species response variable as *S_norm,i_* = *S_i_* / *S_max_*, where *S_max_* is the maximum value of *S_i_* found in any of the landscapes in each of the Figures 1, 3 and 5 (a different *S_max_* value for each figure is considered). The maximum *S_max_* is found in the habitat site that, in each of these figures, has the largest habitat area in the local landscape. In Figures 1 and 5 there are some habitat patches that are larger than the extent of the local landscape (larger than a circle with radius D); hence some of the habitat sites have their local landscapes fully covered with habitat. In the landscapes in Figure 3 this is not however the case, and the habitat site with the maximum values of *A_i_* and *S_i_* has about one fifth of the local landscape covered by habitat. The *S_norm,i_* values are independent from *k* and are easier to interpret, and are hence those shown in Figures S3.1-S3.6: *S_norm,i_*=100% corresponds to the maximum *S_i_* value found in the considered landscapes and 0% to the case of no habitat in the local landscape.

The key parameter in the SARs, as well as in the discussions on the HAH and on the ways to test it (see main text), is the slope parameter *z*, which will drive the specific normalized values of the response variable (*S_norm,i_*) shown for the illustrative landscapes in the figures below.

Obviously, when *z* = 1 the power function converts into a linear function (without need of any logarithmic transformation) as a particular case. The lower the *z* value, the milder the variation in the response variable (e.g. species richness) with changes in habitat amount. In the extreme case, when *z* = 0 there is no effect of habitat area (habitat amount) on species richness, abundance or occurrence.

Several HAH-related studies have tested the HAH by fitting power-law SARs as a linear function of the log-transformed variables and have found the SAR slope (*z*) to range from 0.18 to 0.5 (Haddad et al., 2017), from 0.4 to 0.58 (Rabelo et al., 2017) and from 0.31 to 0.75 (Bueno & Peres, 2019). These values give an average *z* value of 0.45, which I hence use as one of the two slope values applied in the illustrative calculations below. Other studies not related to the HAH have however reported SAR slopes with an average of 0.321 (Triantis et al., 2012), although with a wide variability depending on the cases, ranging from 0.064 to 1.312 (Triantis et al., 2012). I therefore consider here *z* = 0.32 as the second *z* value in the calculations applied to the illustrative cases below. Triantis et al. (2012) noted that the SAR slope (*z*) was however significantly higher when using data sets spanning just two orders of magnitude of area than for data sets spanning more orders of magnitude of area. This is of interest here because the typical range of areas at which the HAH operates may be closer to the case of less orders of magnitude in the variation of area. In particular, Triantis et al. (2012) showed, in his figure 5, that for data sets spanning just two orders of magnitude of area the mean *z* is 0.438. This value is closer to the first value here considered (*z* = 0.45) than to the second one (*z* = 0.32), which corresponds to all SAR datasets and studies reviewed in Triantis et al. (2012) regardless of the ranges of area used.

**FIGURE S3.1.**
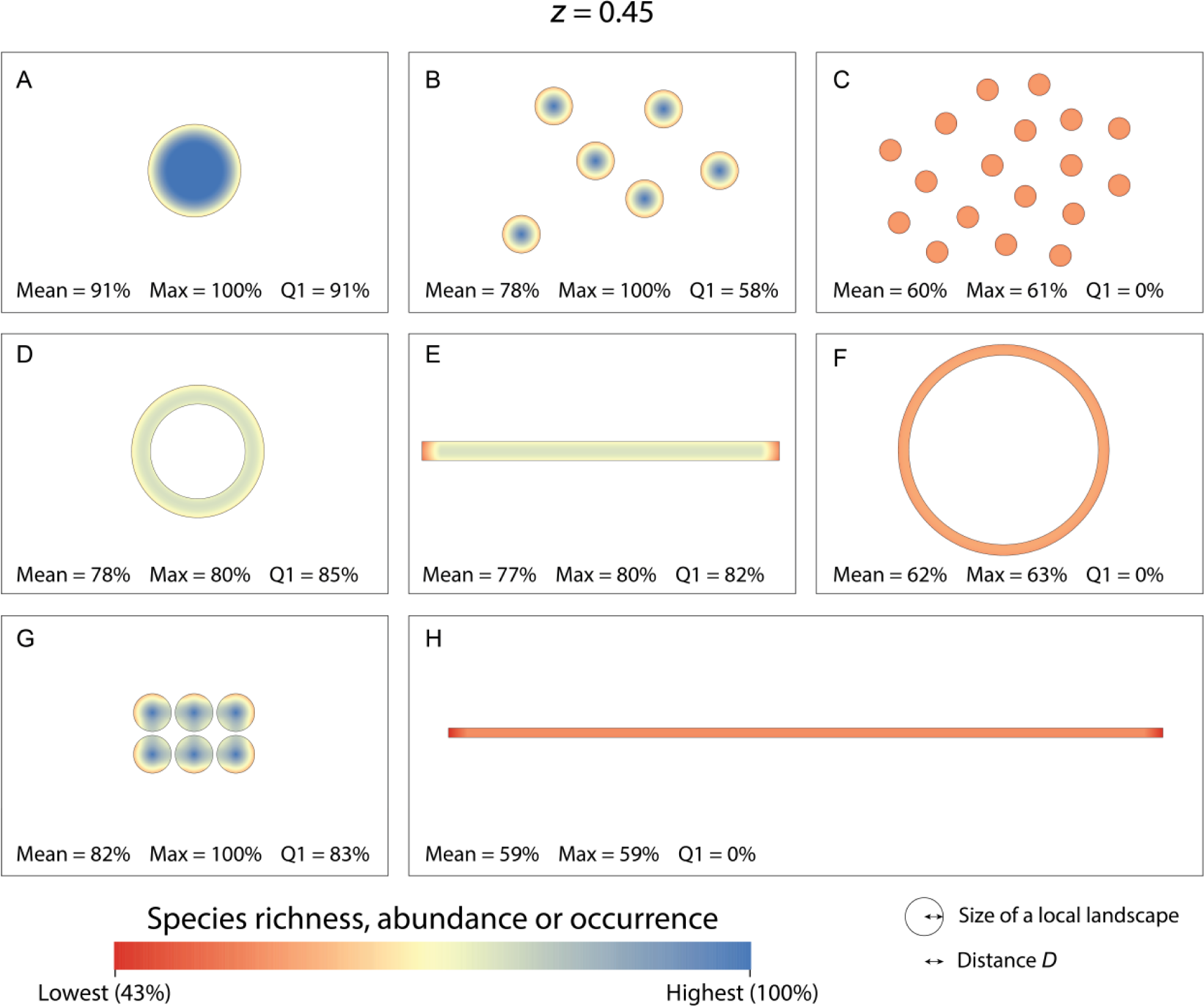
Species richness, abundance or probability of occurrence (*S_norm,i_*) given by the predictions of the Habitat Amount Hypothesis (HAH) in each of the landscapes in Figure 1, calculated as a power law function of the amount of habitat in the local landscape (distance *D*) around each location (habitat site) *i* with the slope parameter (*z*) equal to 0.45. The figure indicates, for each landscape, the mean (Mean) and maximum (Max) value of *S_norm,i_* in the landscape as predicted by the HAH using such power law, as well as the percentage of habitat area in each landscape that has a value of *S_norm,i_* within the top quartile (Q1; within the 75-100% range). The values of the response variable are normalized (*S_norm,i_*) so that 100% corresponds to the maximum value found in these landscapes (which happens when the entire local landscape is covered by habitat) and 0% to the case of no habitat in the local landscape. All habitat sites in these examples have however some amount of habitat in their surrounding local landscapes, so that *S_norm,i_* never goes below 43% in these examples. See Figure 2 in the main text for a similar figure but calculating the response variable as a linear function of the habitat amount in the local landscape; the numbers of Mean, Max and Q1 vary, but they lead to the same trends and conclusions.

**FIGURE S3.2.**
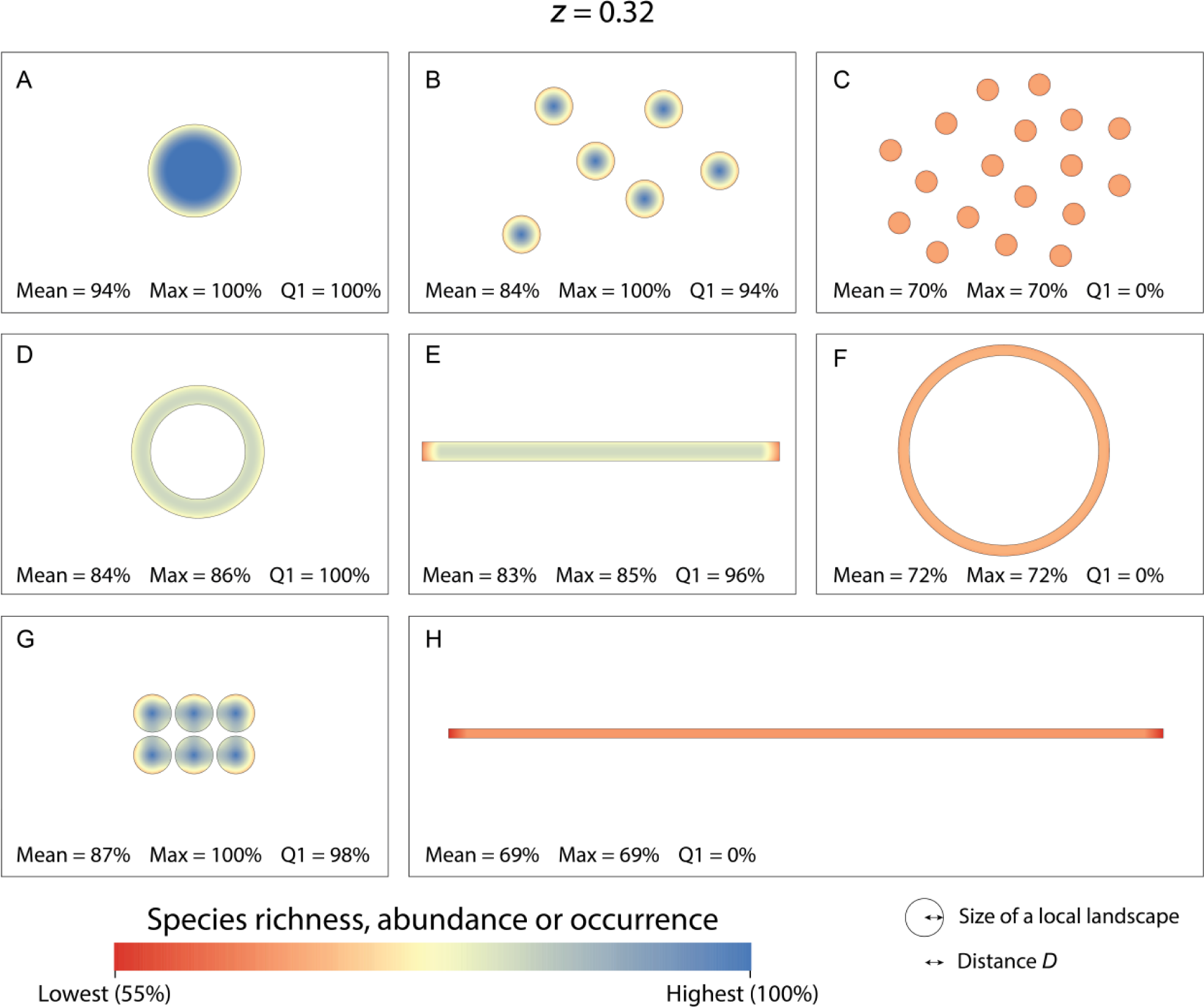
Species richness, abundance or probability of occurrence (*S_norm,i_*) given by the predictions of the Habitat Amount Hypothesis (HAH) in each of the landscapes in Figure 1, calculated as a power law function of the amount of habitat in the local landscape (distance *D*) around each location (habitat site) *i* with the slope parameter (*z*) equal to 0.32. The figure indicates, for each landscape, the mean (Mean) and maximum (Max) value of *S_norm,i_* in the landscape as predicted by the HAH using such power law, as well as the percentage of habitat area in each landscape that has a value of *S_norm,i_* within the top quartile (Q1; within the 75-100% range). The values of the response variable are normalized (*S_norm,i_*) so that 100% corresponds to the maximum value found in these landscapes (which happens when the entire local landscape is covered by habitat) and 0% to the case of no habitat in the local landscape. All habitat sites in these examples have however some amount of habitat in their surrounding local landscapes, so that *S_norm,i_* never goes below 43% in these examples. See Figure 2 in the main text for a similar figure but calculating the response variable as a linear function of the habitat amount in the local landscape; the numbers of Mean, Max and Q1 vary, but they lead to the same trends and conclusions.

### S3.3. Application of the power law to the illustrative landscapes

Figures S3.1 and S3.2 show the results of applying, to the illustrative landscapes in Figure 1 in the main text, the HAH using a power function between the habitat amount in the local landscape and the response variable in the habitat site, with slopes (*z*) equal to 0.45 and 0.32, respectively.

The power law with *z* < 1 produces, as expected, milder decreases in the response variable (e.g. species richness) than the linear relationship (*z* = 1) as a result of spatial configuration changes such as habitat fragmentation (compare Figures S3.1 and S3.2 with Figure 2). These decreases are, as expected, milder for *z* = 0.32 than for *z* = 0.45 (compare Figure S3.2 with Figure S3.1). However, even with such a power-law relationship, the negative effects of habitat fragmentation on the response variable could be made as large as desired by producing increasingly fragmented habitat patterns. Imagine, giving an extreme example, that instead of having the 18 fragmented patches in Figures S3.1C and S3.2C, we would have 600 equally-sized habitat patches arranged similarly as in Figures S3.1C-S3.2C and with the same total area as the single large patch in Figures S3.1A-S3.2A. In such landscape with those 600 patches, the maximum species richness (Max) recorded in the landscape would go down to 13% if using *z* = 0.45 and down to 23% if using *z* = 0.32. These Max values are both below the Max value of 33% in the landscape with 18 patches for the linear case (*z* = 1) in Figure 2C. In any case, regardless of the magnitude of the decline in the response variable with habitat amount and configuration (governed by *z*), and of the specific cases considered, a power law also results, as for the linear case, in the HAH predicting negative effects of habitat fragmentation, elongation, perforation or increased inter-patch distance in all or many of the habitat sites in the illustrative landscapes in Figures S3.1 and S3.2.

Figures S3.3 and S3.4 show the results of applying, to the illustrative landscapes in Figure 3 in the main text, the HAH under a power law relationship between the habitat amount in the local landscape and the response variable in the habitat site, with slopes (*z*) equal to 0.45 and 0.32, respectively.

Similarly to above, the power law with *z* < 1 produces milder decreases in the response variable (e.g. species richness) than the linear relationship (*z* = 1) as a result of spatial configuration changes such as fragmentation, elongation or increased inter-patch distance (compare Figures S3.3 and S3.4 with Figure 4). These decreases are, as expected, milder for *z* = 0.32 than for *z* = 0.45 (compare Figure S3.4 with Figure S3.3). Despite the differences in the magnitude of the decline in the response variable with habitat amount and configuration (governed by *z*), the same trends and general conclusions as for the linear case (see main text) equally hold when using the power law: the Mean values, but not the Max values, decrease with habitat fragmentation, elongation or increased inter-patch distance in the cases illustrated in Figures S3.3 and S3.4.

**FIGURE S3.3.**
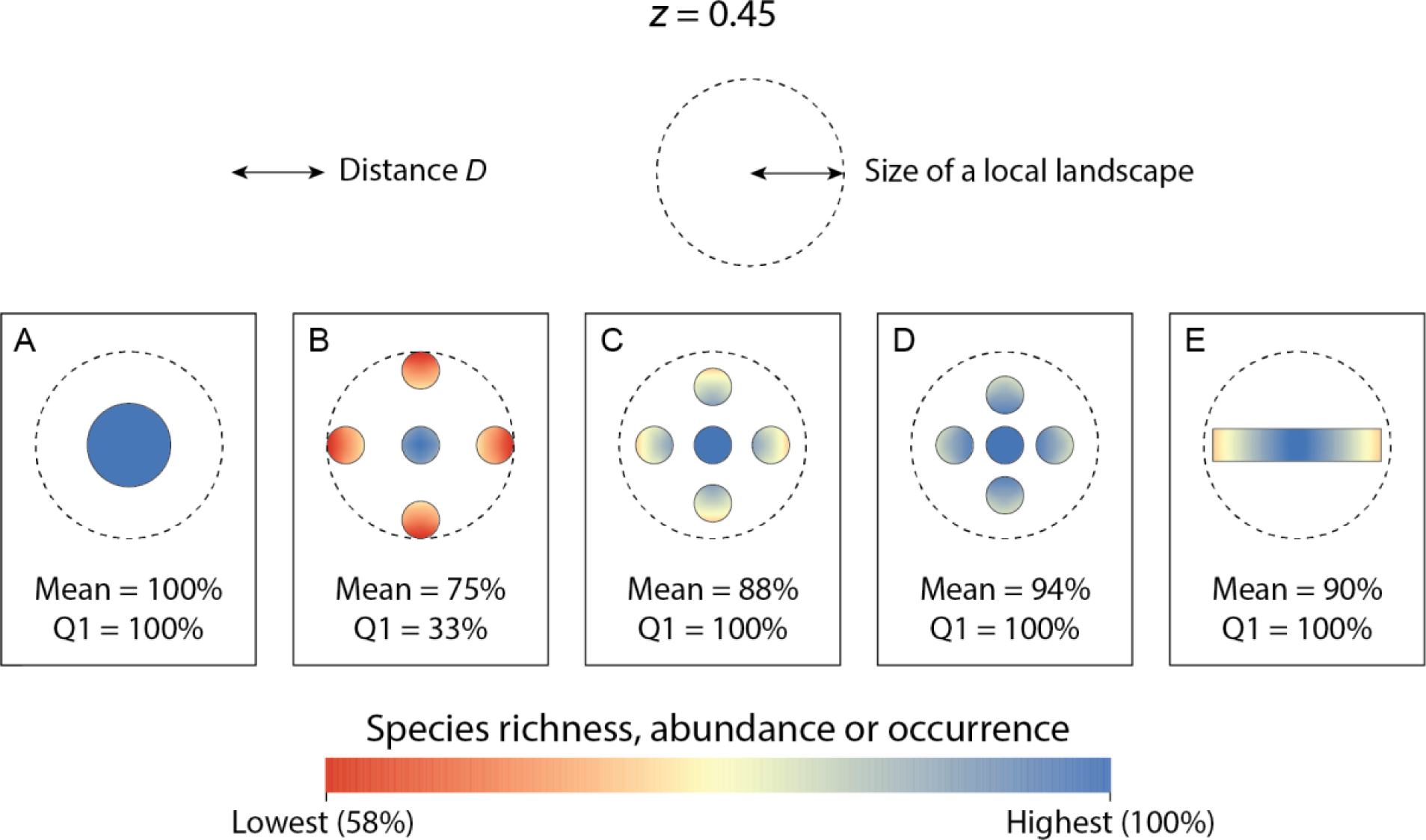
Species richness, abundance or probability of occurrence (*S_norm,i_*) given by the predictions of the Habitat Amount Hypothesis (HAH) in each of the landscapes in Figure 3, calculated as a power law function of the amount of habitat in the local landscape (distance *D*) around each location (habitat site) *i* with the slope parameter (*z*) equal to 0.45. The figure indicates, for each landscape, the mean value (Mean) of *S_norm,i_* in the landscape as predicted by the HAH, as well as the percentage of habitat area in each landscape that has a *S_norm,i_* value within the top quartile (Q1; within the 75-100% range). In all the landscapes there are some habitat sites in which the maximum response variable value is reached (Max = 100%). The values of the response variable are normalized (*S_norm,i_*) so that 100% corresponds to the maximum value found in these landscapes (which corresponds to approximately one fifth of the local landscape covered by habitat) and 0% to the case of no habitat in the local landscape. All habitat sites in these examples have some amount of habitat in their surrounding local landscapes, so that *S_norm,i_* never goes below 58% in these examples. See Figure 4 in the main text for a similar figure but calculating the response variable as a linear function of the habitat amount in the local landscape; the numbers of Mean, Max and Q1 vary, but they lead to the same trends and conclusions.

**FIGURE S3.4.**
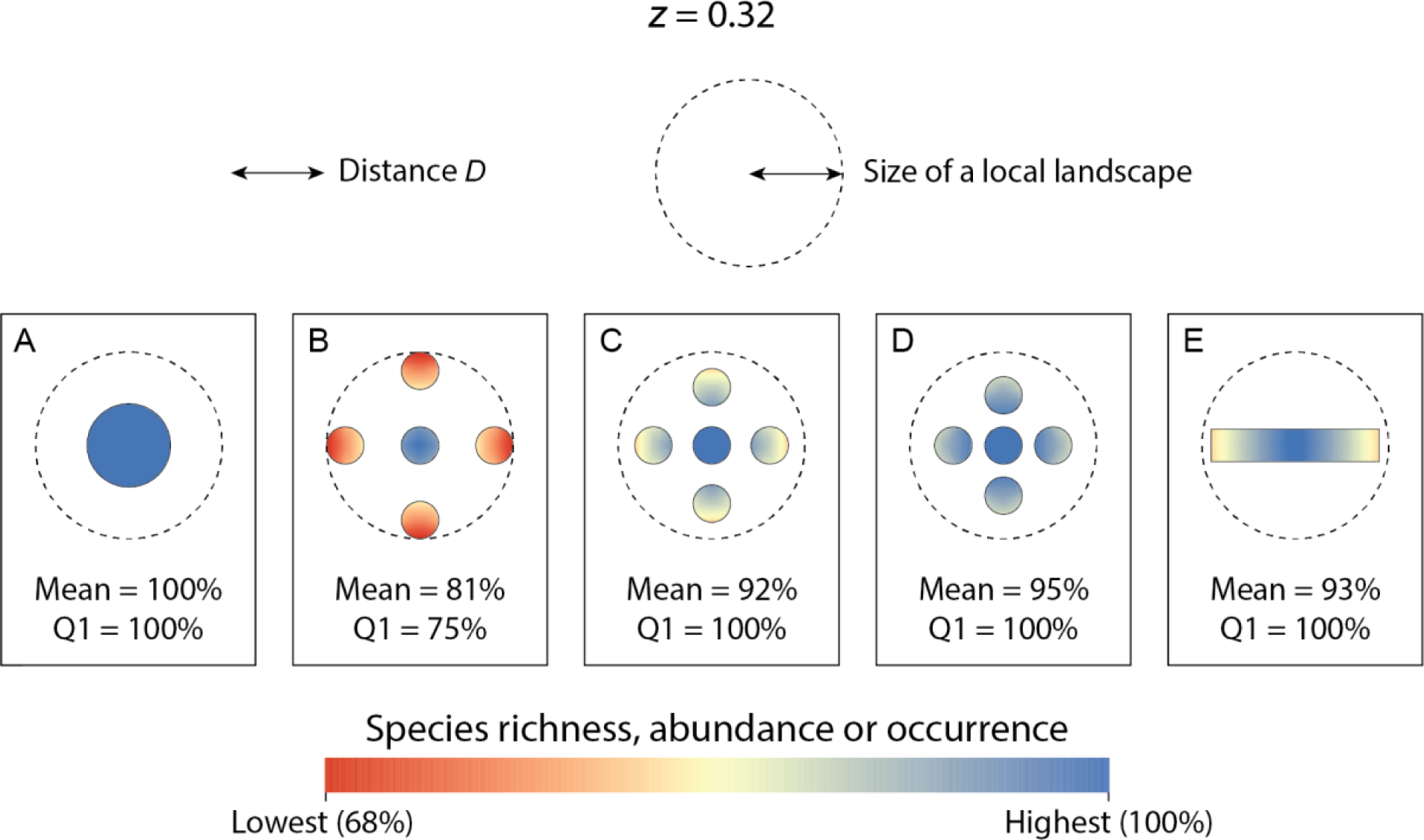
Species richness, abundance or probability of occurrence (*S_norm,i_*) given by the predictions of the Habitat Amount Hypothesis (HAH) in each of the landscapes in Figure 3, calculated as a power law function of the amount of habitat in the local landscape (distance *D*) around each location (habitat site) *i* with the slope parameter (*z*) equal to 0.32. The figure indicates, for each landscape, the mean value (Mean) of *S_norm,i_* in the landscape as predicted by the HAH, as well as the percentage of habitat area in each landscape that has a value of *S_norm,i_* within the top quartile (Q1; within the 75-100% range). In all the landscapes there are some habitat sites in which the maximum response variable value is reached (Max = 100%). The values of the response variable are normalized (*S_norm,i_*) so that 100% corresponds to the maximum value found in these landscapes (which corresponds to approximately one fifth of the local landscape covered by habitat) and 0% to the case of no habitat in the local landscape. All habitat sites in these examples have some amount of habitat in their surrounding local landscapes, so that *S_norm,i_* never goes below 68% in these examples. See Figure 4 in the main text for a similar figure but calculating the response variable as a linear function of the habitat amount in the local landscape; the numbers of Mean, Max and Q1 vary, but they lead to the same trends and conclusions.

Figures S3.5 and S3.6 show the results of applying the HAH to the illustrative landscapes in Figure 5 in the main text, using a power law for the relationship between the habitat amount in the local landscape and the response variable in the habitat site, with slopes (*z*) equal to 0.45 and 0.32, respectively.

The values of the response variable (as summarized by Mean, Max, Q1) are in Figures S3.5 and S3.6 different from those in Figure 6. There are smaller differences between the fragmented patches and continuous habitat in Figures S3.5 and S3.6, and particularly in Figure S3.6, compared to Figure 6. This is a consequence of the lower value of *z* used in each case, which governs the rate of decline of species richness, abundance or occurrence as habitat amount in the local landscape decreases: *z* = 0.32 in Figure S3.6, *z* = 0.45 in Figure S3.5 and *z* = 1 in Figure 6 in the main text. In any case, the same patterns and conclusions as described in the main text and illustrated in Figure 6 hold when considering the power law as in Figures S3.5 and S3.6: the species richness patterns strongly differ between habitat patches and samples areas with the same size in continuous habitat (Figures S3.5 and S3.6). This difference can be such that both mean and maximum species richness decrease in the fragmented habitat patches compared to the equivalent sample areas in continuous habitat (compare Figures S3.5Bf and S3.6Bc, and Figures S3.6Bf and S3.6Bc). Given the difference in species richness in the different-sized patches (Figures S3.5Af and S3.5Bf, Figures S3.6Af and S3.6Bf) and the lack of any difference in species richness in the comparable sample areas in continuous habitat (Figures S3.5Ac and S3.5Bc, Figures S3.6Ac and S3.6Bc), all of which result from the HAH predictions through the power law, it follows that the SAR slope for the fragmented patches has to be steeper than for the continuous habitat, as reported as well in the main text for the linear case.

Therefore, in summary, the conclusions in the main text equally hold when considering a power law: the HAH is compatible with declines in species richness, abundance or occurrence as a result of habitat fragmentation and of other habitat configuration changes, with higher biodiversity for a single large patch than for several small patches with the same total area (SLOSS), and with steeper SAR slopes for more fragmented habitat.

**FIGURE S3.5.**
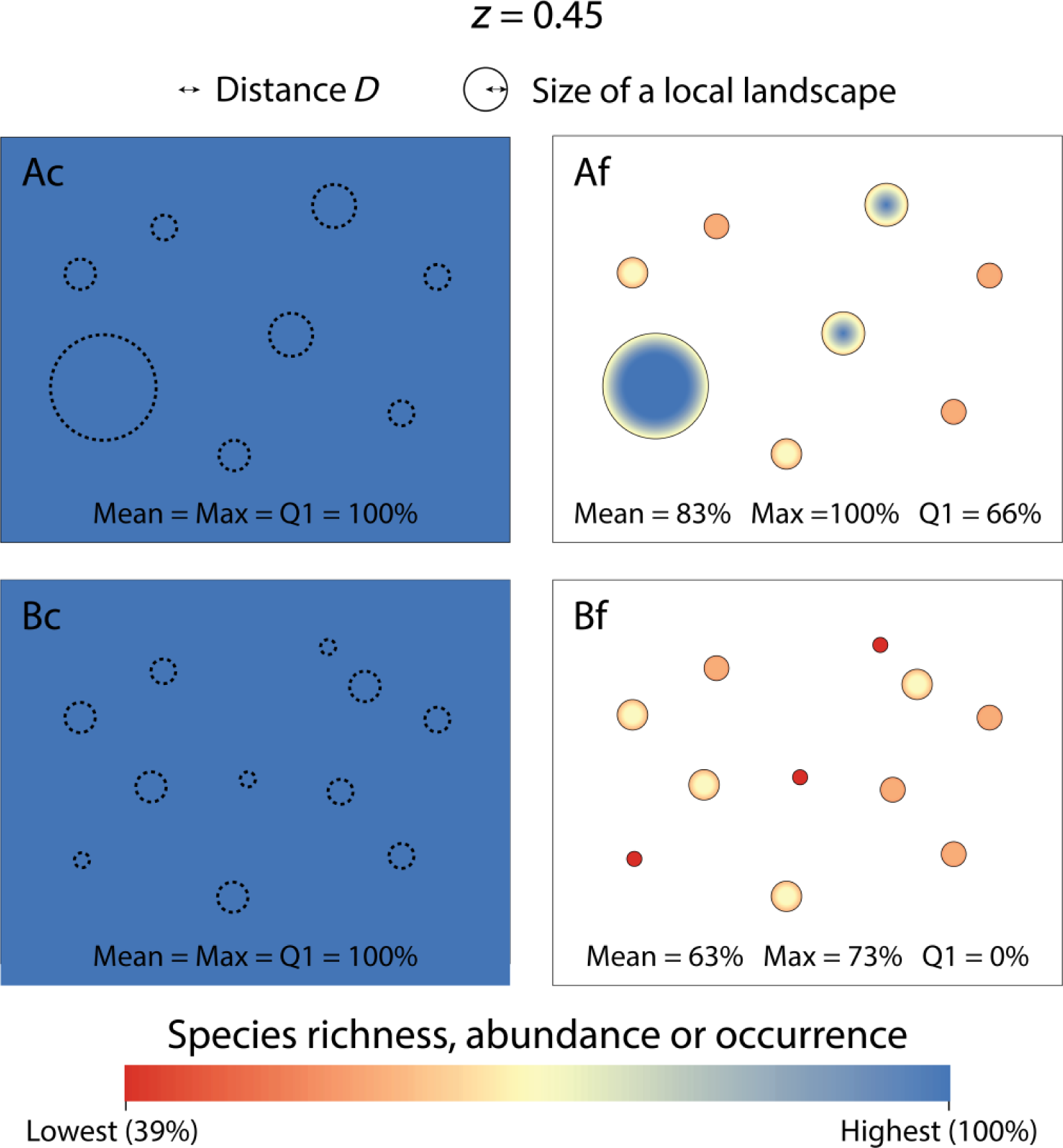
Species richness, abundance or probability of occurrence (*S_norm,i_*) given by the predictions of the Habitat Amount Hypothesis (HAH) in each of the landscapes in Figure 5, calculated as a power law function of the amount of habitat in the local landscape (distance *D*) around each location (habitat site) *i* with the slope parameter (*z*) equal to 0.45. The figure indicates, for each landscape, the mean (Mean) and maximum (Max) value of *S_norm,i_* in the landscape as predicted by the HAH, as well as the percentage of habitat area in each landscape that has a *S_norm,i_* value within the top quartile (Q1; within the 75-100% range). In the landscapes with continuous habitat (Ac and Bc), the habitat is assumed to extend beyond the boundaries of the shown rectangles, and hence the response variable remains high even close to the rectangle border. The values of the response variable are normalized (*S_norm,i_*) so that 100% corresponds to the maximum value found in these landscapes (which happens when the entire local landscape is covered by habitat) and 0% to the case of no habitat in the local landscape. All habitat sites in these examples have however some amount of habitat in their surrounding local landscapes, so that *S_norm,i_* never goes below 39% in these examples. See Figure 6 in the main text for a similar figure but calculating the response variable as a linear function of the habitat amount in the local landscape; the numbers of Mean, Max and Q1 vary, but they lead to the same trends and conclusions.

**FIGURE S3.6.**
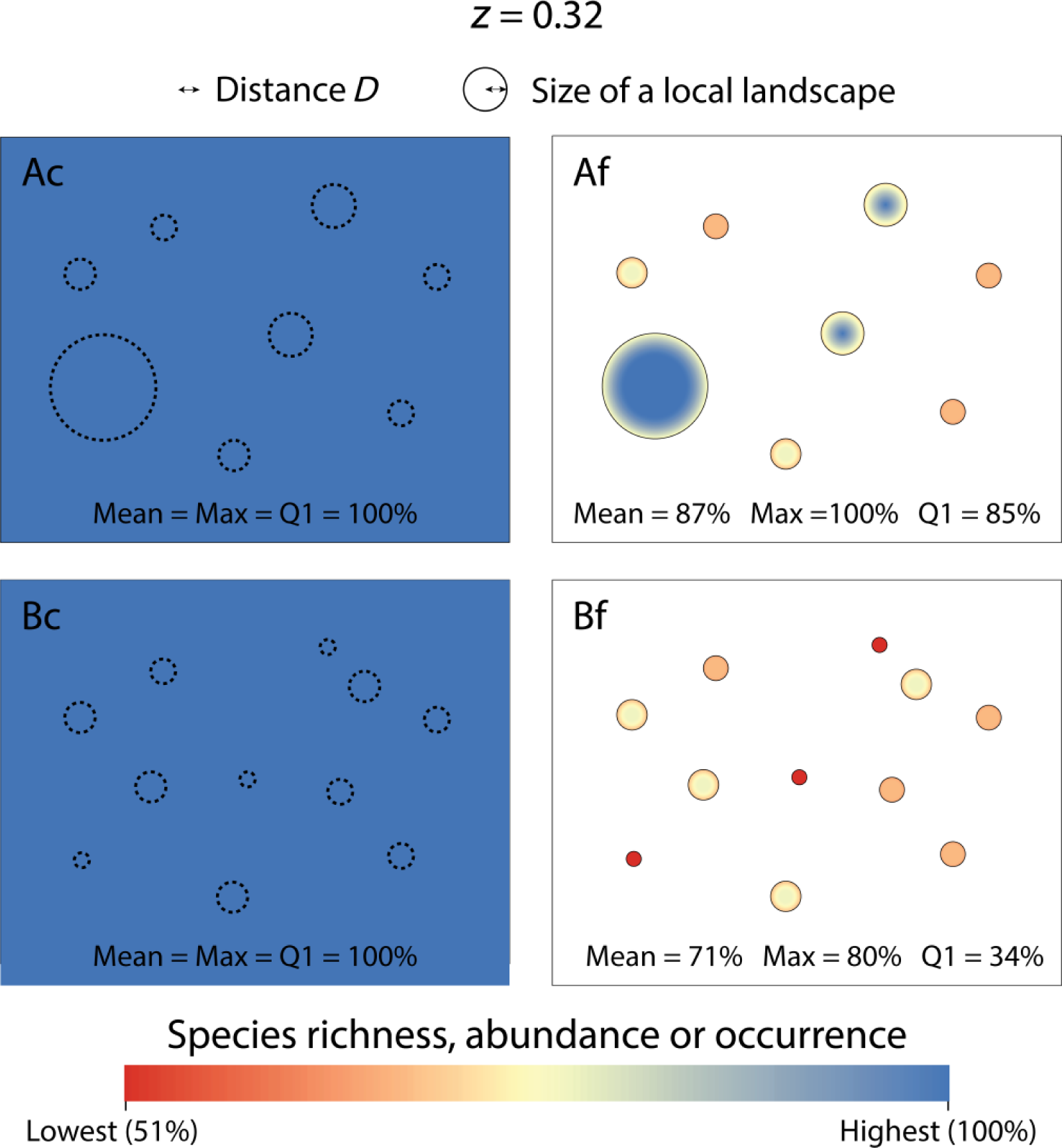
Species richness, abundance or probability of occurrence (*S_norm,i_*) given by the predictions of the Habitat Amount Hypothesis (HAH) in each of the landscapes in Figure 5, calculated as a power law function of the amount of habitat in the local landscape (distance *D*) around each location (habitat site) *i* with the slope parameter (*z*) equal to 0.32. The figure indicates, for each landscape, the mean (Mean) and maximum (Max) value of *S_norm,i_* in the landscape as predicted by the HAH, as well as the percentage of habitat area in each landscape that has a value of *S_norm,i_* within the top quartile (Q1; within the 75-100% range). In the landscapes with continuous habitat (Ac and Bc), the habitat is assumed to extend beyond the boundaries of the shown rectangles, and hence the response variable remains high even close to the rectangle border. The values of the response variable are normalized (*S_norm,i_*) so that 100% corresponds to the maximum value found in these landscapes (which happens when the entire local landscape is covered by habitat) and 0% to the case of no habitat in the local landscape. All habitat sites in these examples have however some amount of habitat in their surrounding local landscapes, so that *S_norm,i_* never goes below 51 in these examples. See Figure 6 in the main text for a similar figure but calculating the response variable as a linear function of the habitat amount in the local landscape; the numbers of Mean, Max and Q1 vary, but they lead to the same trends and conclusions.

## APPENDIX S4. Additional examples of landscapes with varying degrees of habitat fragmentation and all habitat patches smaller than the extent of a local landscape

Landscapes A-C in Figures 1 and 2 in the main text give examples of landscapes with increasing habitat fragmentation (from A to C) while holding constant the amount of habitat in the landscape. The patches are smaller than the extent of a local landscape in landscape C, of the same size than the extent of a local landscape in landscape B, and larger than the extent of a local landscape in landscape A (Figures 1 and 2). Here, I consider in Figure S4.1 a wider set of landscapes with different degrees of habitat fragmentation for the same total amount of habitat, including five cases (landscapes B-F in Figure S4.1) of different patch sizes all smaller than the extent of a local landscape. I apply the HAH predictions to each of the habitat sites in these landscapes, following the same procedure described in section 2.2 (main text) and a linear relationship between the amount of habitat in the local landscape and the response variable in the habitat site, to obtain the results shown in Figure S4.2. These results in Figure S4.2 show that it is possible to decrease the mean and maximum value of the response variable (species richness, abundance or occurrence) as much as desired by producing habitat configurations that are even more fragmented than the one in Figure 2C in the main text.

**FIGURE S4.1.**
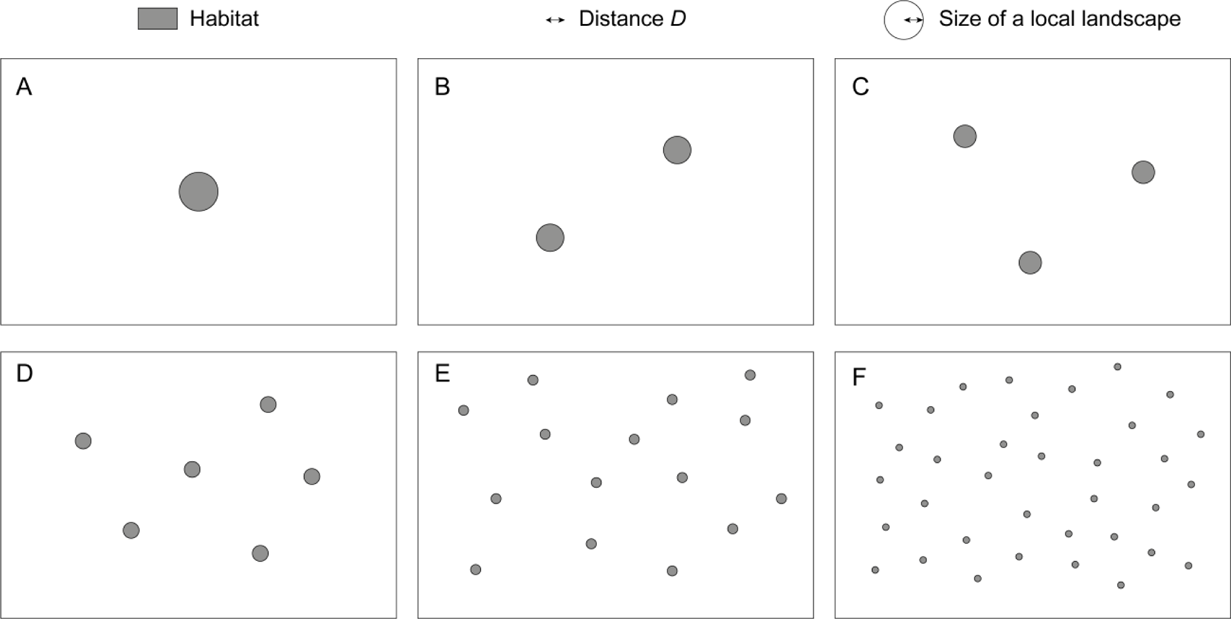
Six landscapes with the same habitat amount (total habitat area) but different degrees of habitat fragmentation, which increases from A to F. All habitat patches in landscapes B-F are smaller than the scale of effect (size of a local landscape). The single patch in landscape A has exactly the same size as a local landscape (as for the patches in landscape B in Figure 1 in the main text). The patches in landscape C in this figure have the same size as the patches in landscape C in Figure 1 in the main text.

**FIGURE S4.2.**
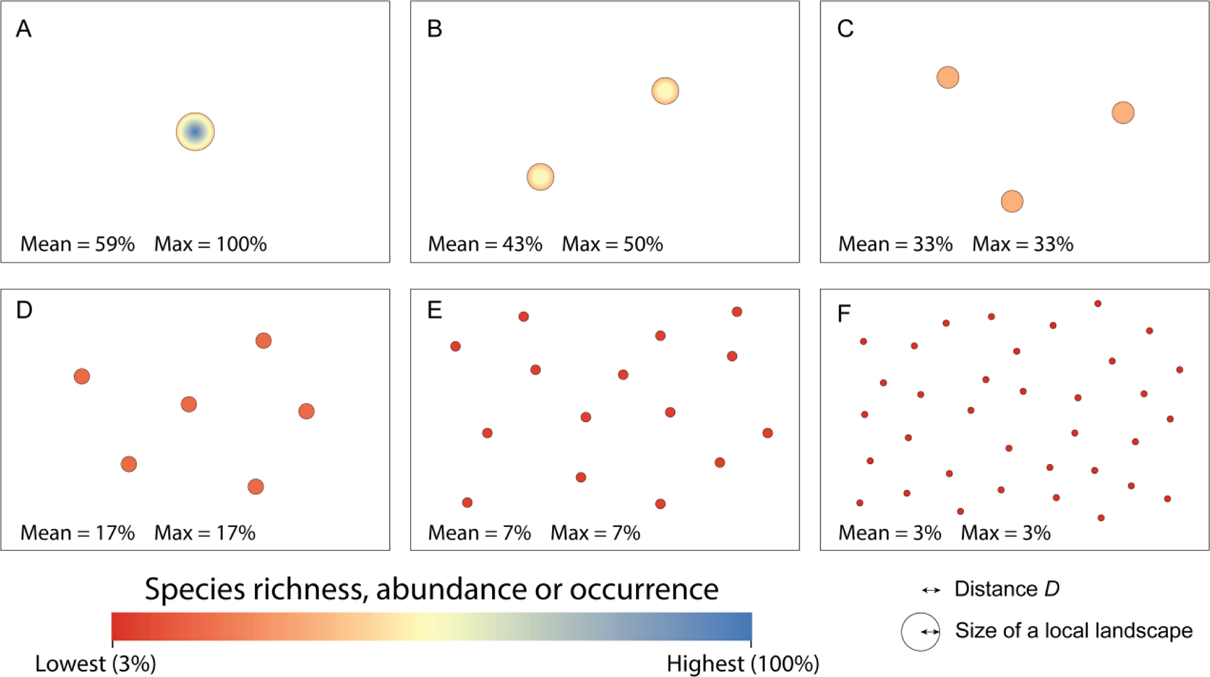
Species richness, abundance or probability of occurrence (*S_norm,i_*) given by the predictions of the Habitat Amount Hypothesis (HAH) in each of the landscapes in Figure S4.1, i.e. calculated for each location (habitat site) *i* as a linear function of the amount of habitat in the local landscape (distance *D*) around the site, as described in section 2.2. The figure indicates, for each landscape, the mean (Mean) and maximum (Max) value of *S_norm,i_* in the habitat sites in the landscape as predicted by the HAH. The results show that an increased habitat fragmentation (breaking apart of habitat) while holding constant the amount of habitat, has negative effects on the species richness, abundance or occurrence in the landscape according to the HAH predictions. The values of the response variable are normalized (*S_norm,i_*) so that 100% corresponds to the maximum value found in these landscapes (which happens when the entire local landscape is covered by habitat) and 0% to the case of no habitat in the local landscape. All habitat sites in these examples have however some amount of habitat in their surrounding local landscapes, so that *S_norm,i_* never goes below 3% in these examples. The values of the percentage of habitat area (habitat sites) in each landscape that has a *S_norm,i_* value within the top quartile (Q1; within the 75-100% range) are not shown in this figure because Q1 is equal to 0% in all cases expect in landscape A, in which Q1=16% (as in landscape B in Figure 2 in the main text).

## APPENDIX S5. Species turnover between habitat sites and the larger-scale predictions derived from the Habitat Amount Hypothesis

The Habitat Amount Hypothesis (HAH) gives species richness (total number of species) in a habitat site, but obviously not the identity of each of the species in the site. The same applies to the HAH-based predictions in Figures 2, 4 and 6 in the main text. When considering an area larger than a single habitat site, such as a habitat patch or an entire landscape comprising multiple habitat sites, the species richness at this larger scale (γ diversity) will be determined both by the species richness at the individual habitat sites (α diversity) and by species turnover between sites (β diversity). While the HAH gives predictions on α diversity, it makes no statement nor gives any information on species turnover (β), i.e. on potential changes, within a considered area, in species composition between different habitat sites with the same habitat amount.

Therefore, with the information provided by the HAH alone it is not possible to directly obtain species richness at spatial scales larger than the habitat site, unless some prior knowledge or assumption on species turnover is made in combination with the HAH. Species turnover can result from a number of factors, such as differences in environmental parameters like climate, species interactions, demographic stochasticity or other factors, which I will here call ‘habitat-independent’ factors. Two habitat sites with the same habitat quality and amount in their surrounding local landscapes will have, according to the HAH, the same species richness, but whether the identity of species in the two sites is the same will depend also on those other habitat-independent factors that may drive species turnover. In any case, the focus here is on the effects of habitat distribution (amount and configuration) on species richness and occurrence, as considered by the HAH, and not on these other habitat-independent factors driving species presence. I therefore here assume that those other habitat-independent factors that may lead to species turnover in the sites do not occur, or put them out of the discussion here since they are out of the scope of the HAH and its predictions. Therefore, I here focus only in the part of the species occurrence that can be explained and predicted by the HAH.

It is therefore appropriate to assume, for the HAH-related purposes in this discussion, that all habitat sites with the same habitat amount in their local landscapes have not only the same species richness, but also the same identity of the individual species found in those sites. It is also appropriate to assume that no new species are found in a given habitat site when the habitat amount in the local landscape around the site decreases. Indeed, if the HAH holds for species richness it is because it holds for the abundance or occurrence of the individual species making up the total number (Fahrig, 2013). If an individual species is only present in a site when the habitat amount is above a given species-specific minimum value, which is the only occurrence-related factor that the HAH can evaluate, it follows that no new species can appear in a site when decreasing habitat amount. For the same reason, an increase in habitat amount can trigger the presence of new species that were not found when habitat amount was lower, but cannot cause the disappearance of species that already existed for lower habitat amounts. If a species would disappear in this case, it would be because of some other habitat-independent factors (e.g. species interactions) that are not considered here since they do not relate to the HAH. The species would not disappear because of the species requirements for a sufficient amount of habitat in the local landscape not being met. I therefore here assume no species turnover between sites with the same habitat amount and nestedness in species composition, i.e. that species assemblages in sites with lower habitat amount are a subset of the assemblages in sites with higher habitat amount in the local landscape.

Under these assumptions, total species richness in an area (e.g. a patch or landscape) comprising multiple habitat sites would be equal to the maximum species richness found in any of the sites found within that area. Therefore, the Max values in Figures 2, 4 and 6 would give the species richness expected in each of the landscapes in this case. This has several clear implications. First, that the HAH is compatible with a decline in the total species richness in the landscape when habitat fragmentation, elongation, perforation and inter-patch distance increase, while holding constant the amount of habitat in the landscape (Figures 2 and 4). For example, total species richness in the landscape (Max value) will decrease from 100% to 33% when a single large habitat patch (Figure 2A) is fragmented into 18 smaller habitat patches with the same total area (Figure 2C). Second, that the HAH is not necessarily SLOSS-neutral, i.e. that it is compatible with higher species richness for a single large patch than for several small patches with the same total area, as given by the same example in Figures 2A and 2C just mentioned. Under the same above-mentioned assumptions, it is also clear that the HAH can also lead, in some other cases, to SLOSS neutrality; consider for examples landscapes Figure 2A and 2B, which have the same Max value. Therefore, whether the HAH is SLOSS neutral or not will depend on the specific cases being considered, and cannot be regarded as a general prediction of the HAH. Third, that the HAH is compatible with steeper SAR slopes for fragmented habitat patches than for sample areas of equal size but contained in continuous habitat. This is clearly apparent by for example considering Figure 6Af, in which the smallest patches have much lower species richness (the response variable is below 35% in all the sites) than the largest ones (in which the response variable reaches 100% in part of the sites), while there is no difference in the HAH-related species richness in the sample areas in Figure 6Ac. Therefore, the slope of a SAR of Type IV (Scheiner, 2003), in which each data point is from a sample corresponding to a unique area (patch), has to be necessarily steeper for Figure 6Af than for Figure 6Ac. The same would hold for other SAR types, such as those constructed by estimating the mean diversity for a given area from multiple combinations of habitat sites.

Whatever departs from these species richness patterns and assumptions would be unrelated to the HAH domain and outside the scope of its habitat amount-based predictions, and would be due to other habitat-independent species turnover and non-nestedness factors. Other more complex assumptions or scenarios on species turnover may complicate the reasoning and the specific numbers in the examples above. It would be however very difficult and infrequent, if not impossible, that in those other assumptions or scenarios species turnover acts in such a way that it always completely compensates, in all cases, the patterns reported above that make the HAH-predictions depend on habitat spatial configuration in the landscape, not be always SLOSS neutral, and lead to SAR slopes that increase with habitat fragmentation.

